# *Atta* Leafcutter Ants are Fine-Scale Bioindicators of Geographic and Seasonal Climate Changes Across the Americas

**DOI:** 10.64898/2026.08.05.743096

**Authors:** Ulrich G. Mueller, Daniela Mera-Rodríguez, Tristan D. Kubik, Tobias G. Mueller, Fabian C. Salgado-Roa, Shreya Rajhans, Ritika V. Bhalla, Madison N. Babb, Rachael Easler, Keiran Irwin-Leventhal, Alejandro G. Di Giacomo, Miguel Vásquez-Bolaños, James Montoya-Lerma, Rainer Wirth, Julián A. Sabattini, Andre Rodrigues, Boris Baer, Francisco Serna, Erika V. Vergara-Navarro, Christian Rabeling, Zachary I. Phillips, Conor A.F McMahon, Sabrina Amador-Vargas, Jon N. Seal, Katrin Kellner, Inara R. Leal, Heraldo L. Vasconcelos, Roberto Camargo, Luiz C. Forti, Jean-Michel Maes, Odair C. Bueno, Martin Bollazzi, Nilson S. Nagamoto, Hermógenes Fernández-Marín, Miguel Limachi, Ronald Zanetti, Fernanda M. P. Oliveira, Maurício Bacci, Cintia M. Santos Bezerra, Martha L. Baena, Richard I. Samuels, Denise D.O. Moreira, Simon L. Elliot, Sergio Sánchez-Peña, Jeffrey Sosa-Calvo, Rachelle M.M. Adams, Jacques H.C. Delabie, Iasmim D. S. Queiroz, John E. Lattke, Orlando Aguilera-Espinosa, Danon C. Cardoso, H. David Hernandez, Flavio Roces

## Abstract

**Aim:** We develop *Atta* leafcutter-ants as bioindicators that respond at fine scales to geographic and seasonal climate changes in the Americas, thereby addressing the paucity of versatile insect bioindicator systems capable of monitoring climate in both Southern and Northern Hemispheres.

**Location:** American tropics and sub-tropics from latitudes S33.6° to N33.2°, with case studies from Colombia, Mexico, and southern USA.

**Time Period:** 2012-2024.

**Taxon Studied:** *Atta* leafcutter-ants.

**Methods:** We elucidate biogeographic patterns of mating-flight phenology of *Atta* leafcutter-ants across the entire *Atta* range from Uruguay/Argentina to the USA, using 2335 records of *Atta* reproductives from the community database iNaturalist, then ground-truth these patterns by comparison with (i) mating-flight records (n=806) from the *Atta* literature; and (ii) mating-flight observations (n=836) accumulated by a consortium of experts who have researched *Atta* for a combined 1000+ work-years. Onset of mating flights can be timed with great precision in *Atta* populations because mass-mating flights are synchronized and triggered by the first major rainfall of a rainy season.

**Results:** Biogeographic patterns in climate-dependent mating-flight phenologies recorded at iNaturalist are corroborated by observations accumulated in the literature and by *Atta* experts. Analyses reveal so-far unknown gradients in mating-flight phenology (e.g., Colombia to USA) that are correlated to geographic climate gradients, and season switches of mating flights from early to late in the year between proximate *Atta* populations (e.g., 200 kilometers apart), for example in the climatically complex Andean regions of Colombia. Regional differences in *Atta* mating-flight phenology correspond to temporal differences in rainfall between ecoregions of Colombia.

**Main Conclusions:** *Atta* ants are tractable insect bioindicators to monitor climate impacts with detailed spatial and temporal resolution across a 9200-kilometer trans-equatorial transect in the Americas. We outline future research directions to explore climate-dependent biology using the continuously growing, and now ground-truthed, information at iNaturalist on *Atta* mating behavior.

## 1 Introduction

Predictive macroecology requires understanding of climate impacts on life-history traits across space and time, yet biogeographic patterns of climate-responsive life-history traits are difficult to elucidate. Most challenging to quantify are ephemeral life-history traits that can be observed only for a short time, such as annual onset of flowering in plants or onset of mating in animals. A key problem here is the logistic challenge of recording observations in many locations at the same time to characterize such life-history traits comprehensively across a large geographic area. For some life-history traits, it is possible to infer biogeographic patterns from museum specimens deposited by collectors over centuries (Helms 2023; Johnson et al. 2023; Qian et al. 2026; GBIF www.gbif.org). Another possibility is to rely on a network of observers who regularly contribute verifiable information to community databases, such as iNaturalist (inaturalist.org) (Feng et al. 2022). Detailed biogeographic patterns can be captured by community databases recording climate-sensitive life-history traits concurrently across vast geographic areas.

Some of the more elusive biogeographic information on arthropods is the spatially variable timing of mating activities (e.g., Kusnezov 1962; Stukalyuk et al. 2022; Helms 2023; Gerlich et al. 2025). In leafcutter ants, for example, mating flights of winged reproductives (alates) of many *Atta* species occur on only a few days each year and, for some species, also only during a short time window of 15-60 minutes in the evening (e.g., *A. vollenweideri*; Fröhle and Roces 2012; Staab and Kleineidamm 2014) or shortly before dawn (e.g., *A. texana* and *mexicana*; Moser 1967; Marti et al. 2015; Mintzer 2018). Despite these observational difficulties, indigenous peoples often command detailed knowledge of regional mating-flight patterns, because *Atta* females are collected as traditional food (Dufour 1987; Costa Neto and Ramos-Elorduy 2006; Aguilera-Espinosa et al. 2024). For example, when one of the co-authors here visited a remote site in Guyana many years ago, a local Amerindian cook impressed the group of visiting myrmecologists with detailed ethnozoological knowledge of leafcutter-ant species, that there exist three different leafcutter species in that area of Guyana (these three valid *Atta* species have different names in the Macushi language), where exactly nests of each of these species can be found, and during which months and at what time of day (e.g., morning, afternoon) each of these *Atta* species have their species-specific mating flights.

In contrast to such detailed ethnozoological knowledge of local mating-flight biology, the literature on *Atta* mating flights is somewhat scattered (Table S4), and there exists no account of biogeographic mating-flight patterns across the entire range of the genus *Atta* from Uruguay/Argentina to the USA. We present here such a biogeographic analysis of *Atta* mating-flight phenologies, using records from the community database iNaturalist, then ground-truth the biogeographic patterns inferred from iNaturalist records by comparisons with published records and with observations accumulated by 48 experts currently researching *Atta* biology. *Atta* reproductives are slow-moving and easy to photograph when returning to the ground after mating flights, and they attract the curiosity of naturalists because of their large size (*Atta* reproductives are 3-4 cm long including wings). The large number of observations of *Atta* reproductives at iNaturalist allows us to quantify variation in mating-flight phenology on spatio-temporal scales with unprecedented resolution. Moreover, because the highly synchronized mass mating flights of *Atta* are triggered by the first major rainfall of a rainy season, and because the onset of a mating-flight season can therefore be timed with great precision in *Atta* populations, we develop *Atta* as a new high-resolution insect indicator of climate, thereby addressing the paucity of versatile insect bioindicator systems capable of tracking climate comprehensively across the Americas (Menéndez 2007; Tiede et al. 2017). Our analyses show that *Atta* mating-flight phenology is a responsive metric of climate shaping biogeographic patterns over (i) short ranges, such as regional differences in *Atta* mating-flight phenology that correspond to temporal differences in rainfall between proximate ecoregions; and (ii) long ranges, such as a 9200-kilometer trans-equatorial transect from latitudes S33.6° (Uruguay) to N33.2° (southern USA) and spanning both Southern and Northern Hemispheres.

## 2 Methods

### 2.1 Data collection

We imported observations of *Atta* reproductives (alate females, dealate females, alate males) deposited at iNaturalist (inaturalist.org) into a customized Excel spreadsheet (Data S1). A detailed account of our methods to curate observations in our dataset is in the Supporting Information (SI). Decisions regarding which observations to include in the dataset were made blind (Kardish et al. 2015) with respect to any biogeographic patterns to be evaluated, and prior to any analyses. Our dataset of 2335 observations covers a ten-year period from 2012 until the end of 2021 (Data S1) and includes observations from 20 countries across the entire *Atta* range from northern Argentina/Uruguay to the southern USA (Table S1).

We included in our dataset all iNaturalist observations that we could verify as *Atta* reproductives, regardless of whether any observation was identified to species, because our aim was to analyze *Atta* mating-flight biogeography at the genus level. Because even expert taxonomists have great difficulty identifying *Atta* reproductives to species, iNaturalist is the most comprehensive repository of observations of *Atta* reproductives, greatly exceeding the number of observations at the Global Biodiversity Information Facility (GBIF https://www.gbif.org/; GBIF requires species identification for each databased record, iNaturalist does not). Although we do not know species identities of most observations in our dataset, it is likely that most of the 15 recognized *Atta* species (Bacci et al. 2009; Barrera et al. 2022) are represented, with the geographically more widespread species (*cephalotes*, *laevigata*, *mexicana*, *sexdens*) likely representing the majority.

### 2.2 Validation and Ground-Truthing

#### 2.2.1 Survey of Mating-Flight Observations by *Atta* Researchers

To assess the validity of the biogeographic patterns inferred from our iNaturalist dataset, we contacted experts currently researching leafcutter-ant biology in locations across the entire *Atta* range. We asked each expert to (i) complete a questionnaire summarizing their personal observations of *Atta* mating flights (Table S3) and (ii) compare their regional observations of *Atta* mating-flights with the biogeographic patterns in mating-flight phenology apparent in the iNaturalist dataset (Figure 1). Detailed methods of our questionnaire survey are in the SI.

**FIGURE 1.**
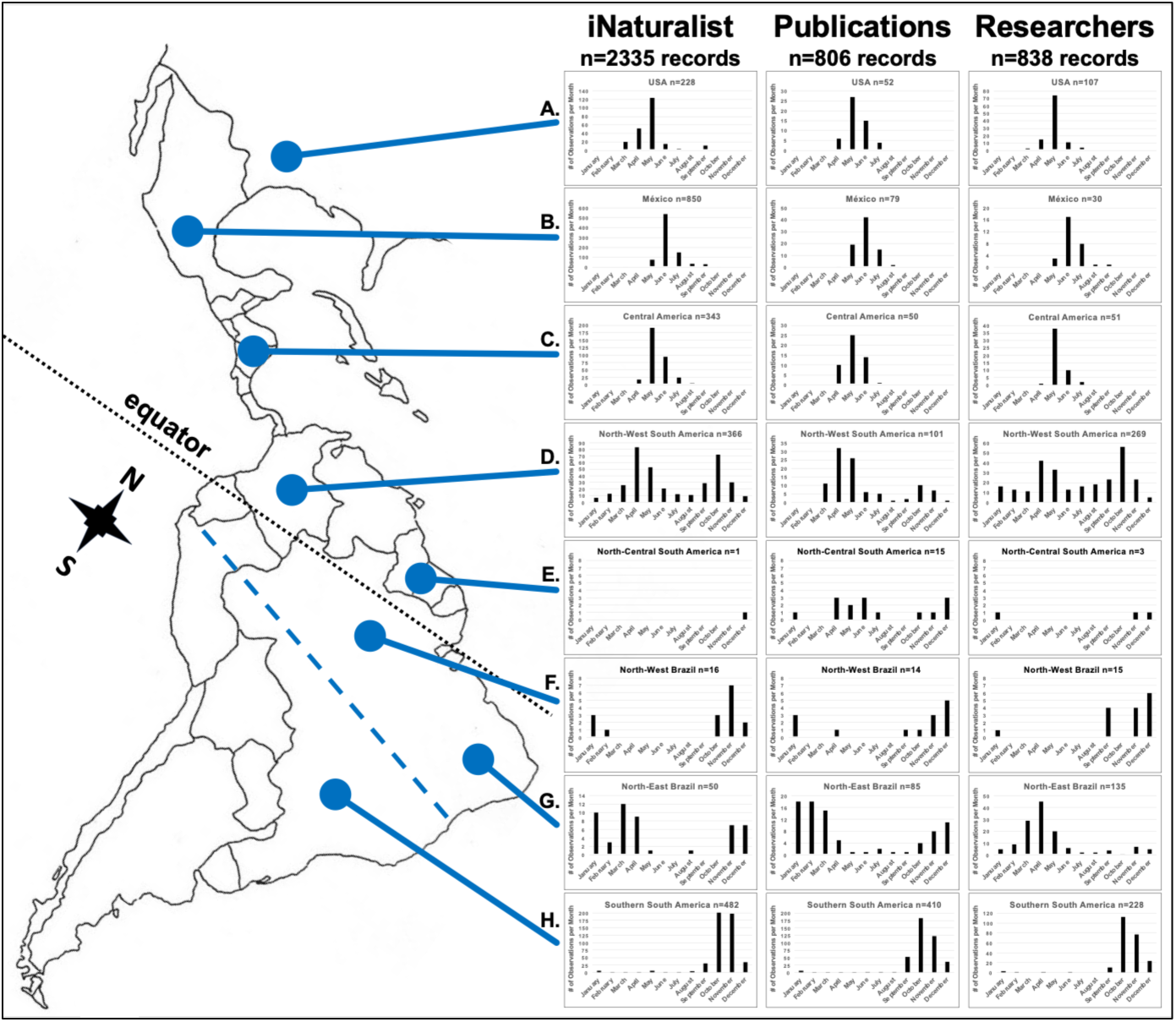
Annual mating-flight phenology of *Atta* leafcutter ants across the entire *Atta* range from Uruguay/Argentina to the southern USA. The leftmost column of graphs summarizes biogeographic patterns inferred from n=2335 records of *Atta* reproductives at iNaturalist from 2012-2021. To ground-truth these patterns, the central column summarizes *Atta* mating-flight records (n=806) reported in the literature (Table S4); the rightmost column summarizes records (n=838) reported in questionnaire surveys of expert *Atta* researchers (Table S3). The graphs summarize regional mating-flight phenology at the genus level, because very few observations can be identified to *Atta* species from iNaturalist photos; this precluded robust species comparisons, except for *A. mexicana* and *A. texana* (Figure 8). **A. USA**, unimodal mating-flight phenology during boreal spring. Almost all observations here are from *Atta texana* (Figures 7&8). **B. Mexico**, unimodal mating-flight phenology during boreal spring. *Atta* mating-flight patterns across México are embedded in a larger mating-flight gradient from north-west Colombia across Central America and México to the southern USA (Figures 5&6). **C. Central America & Cuba**, unimodal mating-flight phenology during boreal spring, combining Guatemala, Belize, Cuba, Honduras, El Salvador, Nicaragua, Costa Rica, and Panamá. **D. North-West South America**, apparently bimodal mating-flight phenology, combining Colombia, Ecuador, northern Peru, Venezuela, Trinidad & Tobago, and the Brazilian States of Roraima and far north-west Amazonas (São Gabriel). Many populations in North-West South America have actually unimodal mating-flight phenology (Figures 3&4), and the apparent bimodality is an artifact of pooling observations across regions where *Atta* have mating flights either predominantly early in the year (April, May) or predominantly late in the year (October, November). **E. North-Central South America** (north of the equator), combining Guyana, French Guiana, Suriname, and the Brazilian State of Amapá. **F. North-West Brazil** (south of the equator), combining the States of Pará and Amazonas (excluding north-west Amazonas). **G. North-East Brazil**, combining the Brazilian States of Maranhão, Tocantins, Piauí, Ceará, Rio Grande do Norte, Paraíba, Pernambuco, Alagoas, Sergipe, and Bahia. **H. Southern South America**, unimodal mating-flight phenology during austral spring, combining Argentina, Uruguay, Paraguay, Bolivia, southern Peru, as well as the Brazilian States of Acre, Rondônia, Mato Grosso, Mato Grosso do Sul, Goiás, Distrito Federal, Minas Gerais, Espírito Santo, Rio de Janeiro, São Paulo, Paraná, Santa Catarina, and Rio Grande do Sul. The dashed blue line bisecting South America is drawn from the southern tip of Ecuador to the northern border of the State of Minas Gerais in Brazil. North of this dashed blue line in the equatorial region of South America, *Atta* mating flights vary regionally; south of this line, *Atta* mating flights peak in October and November. *Atta* mating-flight seasons are prolonged in equatorial latitudes compared to the shorter mating-flight seasons at latitudes distant from the equator.

#### 2.2.2 Survey of Mating-Flight Observations Reported in the Literature

To further validate the biogeographic patterns inferred from our iNaturalist dataset, we completed an exhaustive review of published literature reporting any information on the time of year when *Atta* mating flights occur in specific locations (Table S4). Detailed methods of our literature survey are in the SI.

### 2.3 Colombia

#### 2.3.1 Validation of Mating-Flight Patterns in Colombia, Comparison with Museum Records

To elucidate *Atta* mating-flight patterns across Colombia at greater spatial resolution, we compiled 255 additional records of Colombian *Atta* alates deposited in the Insect Collection of the Universidad Nacional Agronomía Bogotá (UNAB) (Table S3, Questionnaire #07). The UNAB dataset (255 records) and the iNaturalist dataset (260 records) cover many of the same Departamentos (Departments) in Colombia, so we first analyzed annual mating-flight phenology by Department. Because the iNaturalist and UNAB datasets infer similar flight phenologies for different regions of Colombia (Figure S2), we combined for the following ecoregion analyses all Colombia records from iNaturalist and UNAB datasets (Data S4).

#### 2.3.2 Rainfall As Trigger of Mating Flights in Different Ecoregions of Colombia

Because Colombian Departments can span different ecoregions, we characterized annual mating-flight phenology also by ecoregions as defined by Dinerstein et al. (2017). To analyze relationships between mating-flight and rainfall patterns in different ecoregions, we used the rainfall dataset from Urrea et al. (2019). The rainfall dataset and the mating-flight datasets were standardized and paired by day of the year (DOY) (Data S5), as detailed in the SI. We grouped both rainfall and mating flight observations by ecoregions with similar climatic conditions. Using the methods of Curriero et al. (2005), we quantified the temporal relationship between rainfall and mating flights by computing lag-0 correlations, followed by cross-correlation analyses over a ±30-day window to identify the lag with the highest correlation. We tested the correlation between mating-flight and rainfall patterns using RStudio version 4.6.0 (R Core Team 2026) (SI-3 R-Scripts S1&S2).

### 2.4 Analyses of mating-flight phenology of single species *Atta mexicana* and *texana*

We analyzed mating-flight patterns within single species for those regions where only a single *Atta* species occurs, specifically for (i) *Atta mexicana* along a south-to-north transect along the Pacific States of Mexico; and (ii) *Atta texana* along south-to-north and east-to-west transects across the south-central USA. For all other regions, several *Atta* species co-occur that cannot be identified to species from iNaturalist images, so pooling unknown and known *Atta* species within regions enabled us to analyze mating-flight patterns at the genus level.

### 2.5 Expanded dataset for *Atta texana*

To elucidate latitudinal patterns of *Atta texana* mating flights across the south-central USA, we generated an expanded dataset for *A. texana*. We added to our *texana* dataset all subsequent iNaturalist observations of *A. texana* reproductives from the three years 2022-2024. This added 366 more observations of *A. texana* reproductives from 2022-2024 to the 226 *A. texana* observations from 2012-2021, for an expanded dataset of 592 reproductives observed within 13 years (2012-2024).

### 2.6 Expanded dataset for *Atta* in north-east Mexico and Texas

While analyzing latitudinal patterns of *A. texana* mating-flight phenology in the south-central USA (section 2.5), we noticed a marked seasonal switch from mating flights predominantly in August-October in southmost Texas to March-May at all other latitudes across Texas. To understand this shift within a greater latitudinal context, we extended our latitudinal transect across Texas southward to include iNaturalist observations from *Atta* populations in the lowlands along the Gulf of México as far south as Veracruz State. Methods details of designing this 1500-kilometer transect from latitude N20° in Veracruz across Tamaulipas to N33° in northern Texas are summarized in SI-1.

### 2.7 Rainfall Patterns in 2012-2021 Corresponding to iNaturalist Records

To understand how regional rainfall patterns drive *Atta* mating-flight patterns, we obtained monthly total surface precipitation data from the *Precipitation 1.0 degree Data Full Reanalysis* dataset available from the Global Precipitation Climatology Centre (GPCC; https://psl.noaa.gov/data/gridded/data.gpcc.html; Schneider et al. 2022) for the ten years 2012-2021 covered also by our iNaturalist dataset. We imported mean monthly precipitation data into ESRI ArcMaps10.8, projected data in WGS1984, and smoothed data using bilinear interpolation. Details to generate average monthly rainfall maps across the *Atta* range (Figure 2) are in SI-1 Methods. The average rainfall visualized in Figure 2 allowed us to derive expectations regarding corresponding patterns in *Atta* mating-flight phenology.

**FIGURE 2.**
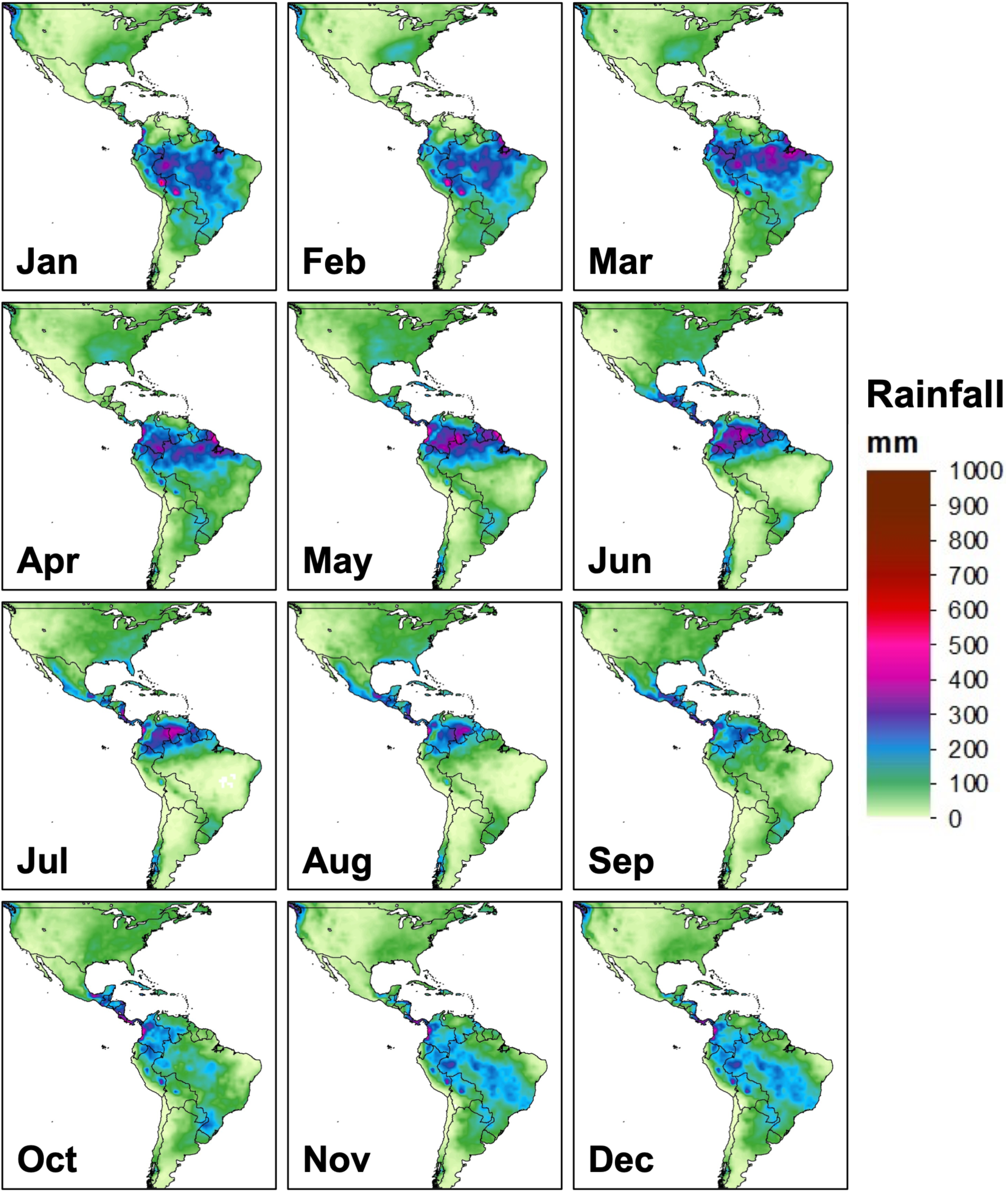
Maps of average monthly surface precipitation (rainfall) across the range of *Atta* leafcutter ants, averaged by month across the ten years 2012-2021 covered by our iNaturalist dataset of *Atta* reproductives. Rainfall data are from the Global Precipitation Climatology Centre. Spatial and temporal variation in rainfall predicts corresponding variation of *Atta* mating flights, visualized in Figures 3-8. Figure S1 shows monthly rainfall for each month from 2012-2021. Supplementary file SI-2 is a GIF animation of these monthly precipitation changes from January 2012 until December 2021.

### 2.8 Correlations Between Julian Observation Dates and GPS Locations

We used R-version 4.4.1 in RStudio (R Core Team 2026; Posit Team 2025) to evaluate statistical support of correlations between latitude (or longitude) and ordinal Julian dates for (i) *Atta* mating flights across Central and North America, and for (ii) *Atta texana* mating flights across the south-central USA. We calculated Spearman correlations because some of the dependencies of mating-flight dates on latitude or longitude were non-linear, and because Julian dates are bounded (between 1-366 days) and thus unlikely normally distributed (SI-3 R-scripts S3&S4).

## 3 Results and Discussion

### 3.1 Mating-Flight Patterns Across the Entire *Atta* Leafcutter-Ant Range

Figure 1 summarizes latitudinal patterns of *Atta* mating flights across the entire *Atta* range from S33.5° southern latitude in Entre Ríos, Argentina, to N33.2° northern latitude in Texas, USA. This analysis summarizes mating-flight phenology at the genus level (i.e., not at the level of individual *Atta* species), because most observations of *Atta* reproductives at iNaturalist are identified only to genus level (Data S1) and consequently sample sizes were very small for most identified *Atta* species (Table S1; SI-1 Footnote 1), precluding robust comparisons between species, except for *A. mexicana* and *A. texana* (3.6 below).

In southern South America, south of the blue dashed line in Figure 1 drawn from southernmost Ecuador at the Pacific coast to the northern border of the State of Minas Gerais in Brazil near the Atlantic coast (SI-1 Footnote 2), *Atta* mating flights occur unimodally during the austral spring from September to December, with peak mating flights in October and November (Figure 1H). This single peak in mating flights in southern South America is a well-known constant in indigenous knowledge, folklore, and myrmecology (e.g., Autuori 1949; Kusnezov 1962; Mariconi 1970; Table S4). Mating-flight patterns in Central America and North America are likewise unimodal, with peaks in May-July during the boreal spring (Figures 1 A-C; Figures 5-8). In southern South America, Central America, and North America, both temperature and rainfall shift between dry season in winter and wet season starting in spring (Figure 2), and the *Atta* mating-flight season starts at that transition from dry to wet season (Tables S3&S4).

In the regions near the equator in South America (hereafter called northern South America, north of the blue dashed line in Figure 1), *Atta* mating-flight seasons extend across many months or even the entire year (Figures 1 D-G), much longer compared to the shorter mating-season windows of 2-3 months in Central America, North America, and southern South America (Figures 1 A-C&H). Two reasons prolong the mating-flight season in northern South America: (a) rainfall patterns in the equatorial regions of South America are less seasonal, particularly in north-west South America (Marengo and Espinoza 2016; Figure 2); and (b) monthly lowland temperatures are more constant, sufficiently warm throughout the year to permit *Atta* flights potentially every month. Long-term or cyclical climate changes may eliminate rainfall continuity throughout the year in north-west South America, possibly creating more pronounced transitions from dryer to wetter seasons in some regions, to which *Atta* may respond with more seasonal mating flights triggered by rains following anomalous dry seasons.

#### 3.1.1 Ground-Truthing iNaturalist Records by Comparison with Observations by *Atta* Experts and with Records from Literature

Biogeographic patterns in *Atta* mating-flight phenologies inferred from 2335 iNaturalist records (Figure 1) were corroborated by (i) 836 mating flights observed by 48 *Atta* experts who completed our questionnaire (Figure 1; Table S3, Data S2) and who have researched *Atta* biology for a combined 1000+ work-years; and (ii) 806 mating flights reported in the *Atta* literature (Figure 1; Table S4, Data S3). As above, we summarize mating-flight phenologies at the genus level because, in the expert and literature datasets, no clear differences emerged when comparing mating-flight phenologies between sympatric *Atta* species (i.e., sympatric *Atta* species have comparable mating-flight phenologies; SI-1 Footnote 1, Table S5); we therefore pool information from different *Atta* species in all below biogeographic analyses. At the genus level, the congruence between mating-flight phenologies inferred from iNaturalist observations, expert observations, and literature records is best for the regional datasets with larger sample sizes (southern South America, North-West South America, Central America, Mexico, USA), somewhat less so for smaller regional datasets (North-Central South America, North-West Brazil, North-East Brazil (Figure 1), suggesting that possible discrepancies between the three sources of observations is a sample-size problem. Because three times more records were accumulated within 10 years (2012-2021) by the iNaturalist community, compared to 1000+ work years by 48 *Atta* experts and to 150 years of *Atta* publications, iNaturalist emerges as the richest source of information (Table S2) to quantify future climate-driven changes in mating-flight phenology of *Atta* ants.

### 3.2 North-West South America

One of the most interesting patterns is an apparently bimodal phenology of *Atta* mating flights in north-west South America, with peaks in April and October, and intervening lows in December/January and July/August (Figure 1D). North-west South America has significant rainfall every month throughout the year (Marengo and Espinoza 2016; Figure 2), and for most ecoregions there is no pronounced dry season as in nearby Central America. *Atta* could respond to this rainfall continuity with at least some mating flights most months throughout the year, but separate fine-grained spatial analyses indicate that *Atta* mating-flight phenology is unimodal in most locations across north-west South America (Figures 3&S2C) and that the apparent bimodality in Figures 1D&S2A is an artifact of pooling observations across heterogeneous regions where mating flights occur either early in the year (April, May) or late in the year (October, November) (Figures 3&S2C). In south-west Colombia, for example, *Atta* mating flights occur almost exclusively in October and November, whereas they occur April-June throughout the rest of Colombia, as they do also in neighboring Panamá, Venezuela, and north-west Brazil north of the equator (Figure 3). When summarizing observations across all of Colombia as in Figures 1D&S2A, therefore, the pronounced bimodality of the mating-flight phenology emerges as an artifact of pooling mating-flight observations across the climatically complex Andes region of Colombia. In Ecuador, mating-flight phenology differs between west and east of the Andes that bisect Ecuador north to south (Figures 3G&H).

**FIGURE 3.**
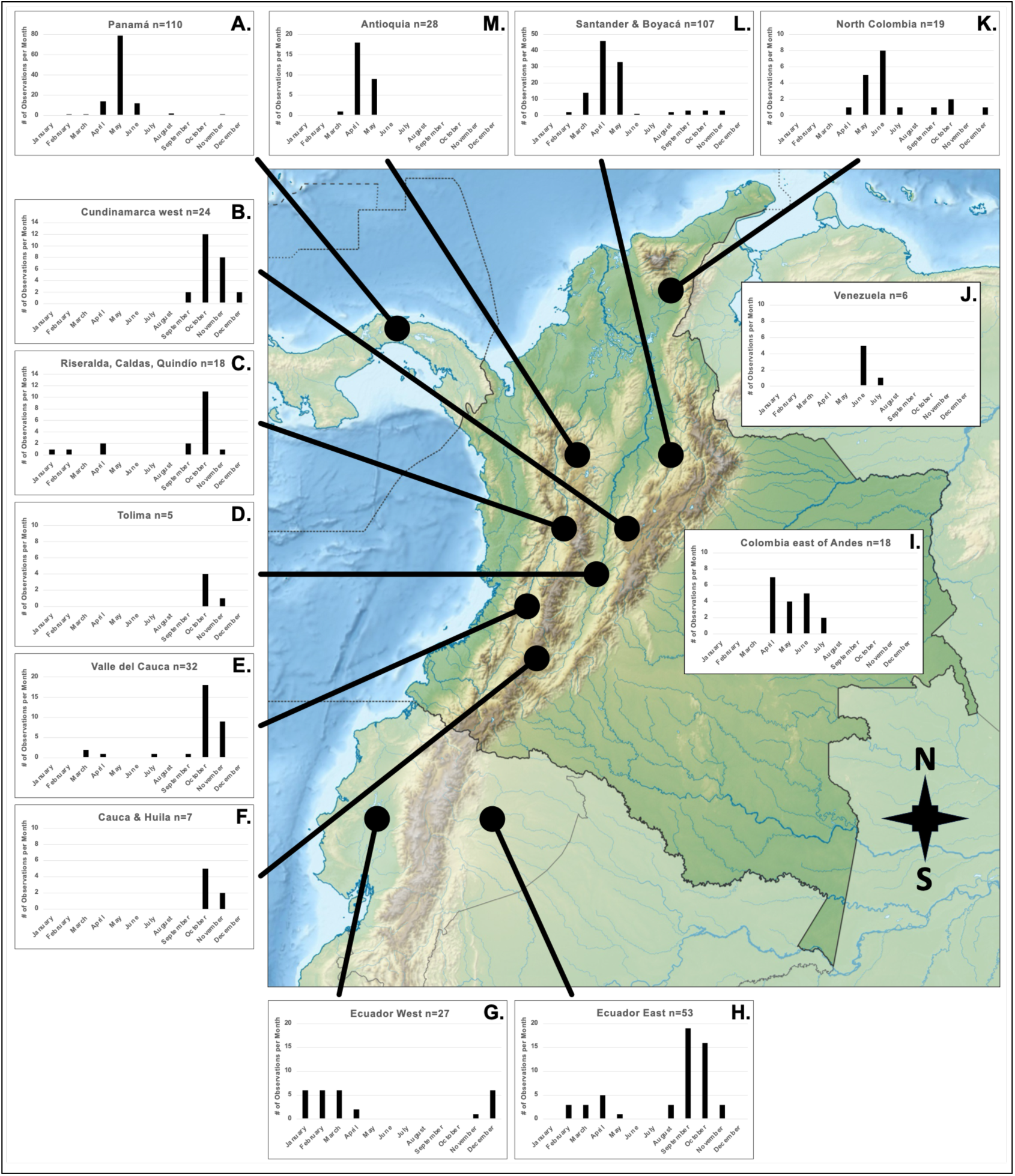
Mating-flight phenology of *Atta* leafcutter ants in Colombia and the neighboring countries Panamá (**A**), Ecuador (**G** & **H**), and Venezuela (**J**). *Atta* has mating flights predominantly in April-July in most of Colombia and likewise in Panamá and Venezuela, whereas *Atta* has mating flights predominantly in October and November in south-west Colombia (**B**-**F**). When pooling observations across all of Colombia and nearby countries, therefore, mating-flight phenology appears bimodal (Figure 1D) as an artifact of pooling across regions with very distinct mating-flight phenologies within Colombia and also within Ecuador. In Ecuador, mating-flight phenology differs between west (**G**) and east (**H**) of the Andes; only east of the Andes in Ecuador appears to be a bimodality in annual mating-flight phenology. In central Colombia, there exists an interesting seasonal switch in mating-flight phenology across a well-sampled region ranging from the Departments of Antioquia (n=26 observations), Santander (n=103) and Boyacá (n=4) to the somewhat more southern Departments of Cundinamarca (western half, west of Bogotá; n=24), Caldas (n=1), Risaralda (n=13), and Quindío (n=4) (see also Figure S2C). While this transect spans only about 200 km north to south, *Atta* mating flights occur predominantly in April/May in Antioquia (**M**) and Santander & Boyacá (**L**), but predominantly in October/November in Cundinamarca (**B**) and Risaralda, Caldas, and Quindío (**C**). This seasonal switch in *Atta* mating-flight phenology across a 200-km north-south transect is likely driven by seasonal rainfall variation interacting with elevational gradients across this Andean region. Relief map from Wikimedia, CC BY-SA 3.0.

#### 3.2.1 Colombia

*Atta* exhibit unimodal mating-flight phenologies with short mating-flight seasons in some Colombian Departmentos (administrative Departments), but prolonged mating-flight seasons or even bimodal mating-flight phenologies on other Departments (Figure 3, Figure S2). For Cali in the Valle de Cauca Department, a bimodal mating-flight phenology was already reported by Montoya-Lerma et al. (2006, 2012), driven there by annual bimodal rainfall patterns (peak rainfall April-May and October-November; Armbrecht et al. 2012). When comparing adjacent Departments, we found a marked transition in mating-flight phenology across a well-sampled region ranging from the Departments of Antioquia and Santander to the somewhat more southern Central Departments including Cundinamarca (western half, west of Bogotá), Caldas, Risaralda, and Quindío (Figure 3, Figure S2C). While this region spans only about 200 km north to south, *Atta* mating flights occurred predominantly in April and May in Antioquia and Santander, but predominantly in October and November in the Central Departments (Figure 3, Figure S2C). In this region of central Colombia, the four species *Atta cephalotes*, *A. colombica*, *A. laevigata*, and *A. sexdens* are sympatric, with *A. cephalotes* being the most abundant (Fernández et al. 2015; Castaño-Quintana et al. 2019). Marked differences in relative abundances of these four different *Atta* species across this 200-km north-south transect therefore do not appear to explain the seasonal switch from spring to fall mating flights within 200 km. Geographic differences in rainfall patterns in Colombia (see below) driving distinct mating-flight phenologies is the more plausible explanation for this short-range switch, rather than geographically-varying abundances of the four *Atta* species combined with species-specific specialization on different flight seasons early or late in the year.

Santander (in Cordillera Oriental) and Antioquia (in Cordillera Central) share somewhat similar climatic patterns although they are on different sides of the Andes, because rainfall is influenced here more by the Caribbean than the Pacific (Espinoza et al. 2020). In contrast, Quindío, Risaralda, Caldas, and the western half of Cundinamarca are in the core of the Cordillera Central, and rainfall patterns there are largely driven by the Pacific (Espinoza et al. 2020). Because the Andes are a dispersal barrier to *Atta* ants beyond isolation-by-distance (Muñoz-Valencia et al. 2022, 2023; Barrera et al. 2022), and because the Andes have been identified as a definitive dispersal barrier to leafcutter-ant fungal cultivars (Mueller et al. 2017), the landscape heterogeneity of Colombia is inherently interesting for understanding the interaction of *Atta* eco-geography, life history and behavior, including the short-range shifts in mating-flight phenology documented here (Figure 3, Figure S2).

Regional differences and short-range transitions in *Atta* mating-flight phenology across Colombia are undoubtedly related to the topography of three Andean Mountain chains crossing in north-south direction (Occidental, Central, and Oriental Cordilleras; Figure 3), interacting with annual northward and southward shifts in the Intertropical Convergence Zone (ITCZ) and with Pacific and Caribbean oceanic influences that drive annual rainfall dynamics in Colombia (Bedoya-Soto et al. 2019). In the coastal regions of northern Colombia, there exists a distinct dry season from December to March (Arregocés et al. 2024) as in nearby Venezuela and Panamá, and the predictable onset of spring rains in April/May trigger *Atta* mating flights in these regions (Figures 3A&K). In eastern Colombia (Llanos Orientales savannah), there exists a single-peak rainy season (Etter and Botero 1990) with an onset of rains in April/May, corresponding to the single-peak *Atta* mating flights observed in eastern Colombia (Figure 3I, Figure S2C). In contrast, the Andean Mountain chains generate local rain shadows such that proximate regions can differ markedly in precipitation patterns in central Colombia (Urrea et al. 2019). For example, an annual bimodal rainfall pattern is well-documented for the Andean region (Poveda et al. 2005; Urrea et al. 2019) and also for specific localities like the Aburrá Valley encompassing the Medellin metropolitan area (Antioquia Department; Bedoya-Soto et al. 2019) where most iNaturalist observations from Antioquia were collected, with the driest season in January and February, followed by two distinct wet seasons in March-May and September-November. *Atta* mating flights in the Medellin metropolitan area occur in April/May (Figure 3M) coincident with the onset of the first wet season, but not in September-November (i.e., the onset of the second wet season does not trigger a second peak of major mating flights). This suggests three hypotheses on interacting climate factors that drive *Atta* mating phenology. First, a sharp transition from a true dry to wet season is the most important cue triggering *Atta* mating flights, as noted also in our Tables S3 and S4. Second, the onset of a dry season could be a potential cue stimulating production of alates in *Atta* nests, because colonies need to start rearing of alate brood about 2-3 months before mating flights and alate development requires about 70-90 days. Third, the second rainfall peak in September-November in the Medellin area does not trigger a second round of mating flights because this second rainfall peak is not preceded by a true dry period stimulating alate production.

#### 3.2.2 Museum Records Verify *Atta* Mating-Flight Patterns in iNaturalist Data from Colombia

When pooling records across Colombia, a bimodal annual flight phenology emerges in the iNaturalist dataset (n=260 records; Figure S2A) and also in a separate dataset compiling records of *Atta* alates deposited in the Insect Collection of the Universidad Nacional Agronomía Bogotá (UNAB) (n=255 records; Figure S2B; data from Questionnaire #07 in Table S3). Both datasets show an April peak and an October peak in mating flights (compare Figures S2A and S2B). Unlike the iNaturalist dataset where most observations cannot be identified to *Atta* species, the UNAB dataset includes mostly records of identified species, allowing species comparisons. Sympatric populations of the four *Atta* species represented in the UNAB dataset have comparable annual mating-flight phenologies (Table S5, SI-1 Footnote 1). At the genus level, there is a clear differentiation between south-west Colombia (*Atta* mating flights occur late in the year) and the rest of Colombia (flights early in the year) (Figures S2C and S2D). This indicates, as already noted in the discussion of Figure 3, that the overall bimodal flight phenology apparent in Figures S2A and S2B is an artifact of pooling data from locations with different flight phenologies. Interestingly, the UNAB data confirm the aforementioned short-range switch (Figure 3) in mating flights from primarily April/May in Santander to primarily October/November only 200 km further south in Cundinamarca (compare Figures S2C and S2D). Because iNaturalist and UNAB datasets infer similar flight phenologies for different regions of Colombia (Figure S2), we combine iNaturalist and UNAB datasets for the following ecoregion and cross-correlation analyses (Data S4, Data S5).

#### 3.2.3 Rainfall Triggers Different Mating-Flight Phenologies in Different Ecoregions of Colombia

The normalized curves of mating-flight patterns and rainfall exhibit similar trends across most ecoregions in Colombia, particularly in well-sampled ecoregions such as Dry Forests and Montane Forests (Figure 4B). In these ecoregions, rainfall and mating-flight activity show closely synchronized seasonal patterns, suggesting a strong relationship between these parameters. However, in ecoregions with fewer observations, such as Llanos and Xeric Scrub ecoregions, the correlation between the two variables is less pronounced (Figure 4B).

**FIGURE 4.**
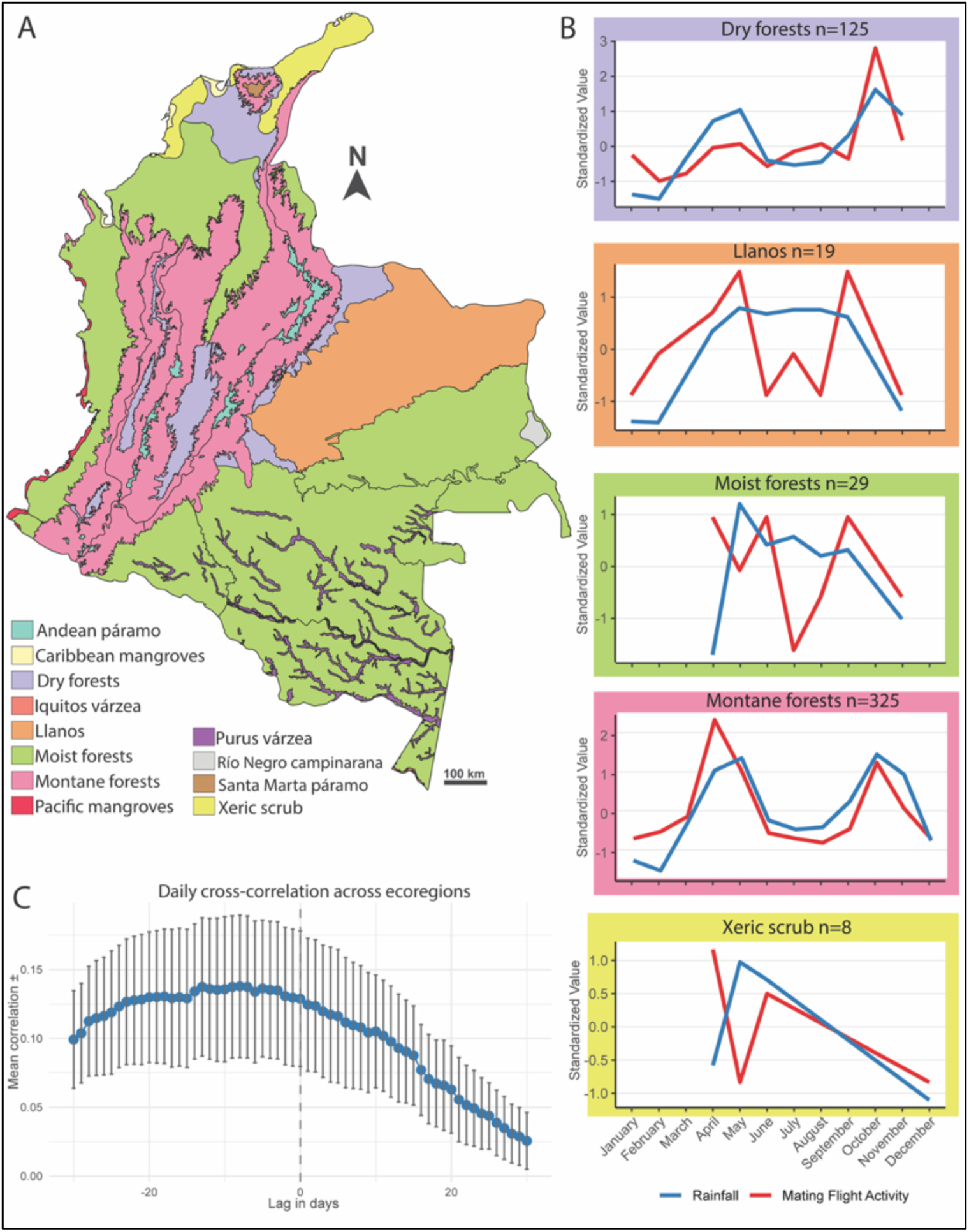
Cross-correlation between rainfall and *Atta* mating-flight activity in Colombia. **A**. Map of Colombian ecoregions as defined by Dinerstein et al. (2017). **B**. Normalized seasonal patterns of rainfall (blue) and mating flight activity (red) across five different ecoregions with sufficient mating-flight observations. We normalized (z-scored) mating-flight observations and rainfall values such that z-scored mating-flight and rainfall values can be projected onto the same y-axis of a single graph. **C**. Daily cross-correlation between rainfall and mating-flight activity across a ±30-day window (mean ±1 SE across six ecoregions with sufficient alate observations). Negative lags indicate that peak mating-flight activity precedes peak rainfall. In ecoregions with sufficient flight observations (Dry and Montane Forests), the peak in mating-flight activity precedes the peak in rainfall by 5-10 days.

#### 3.2.4 Cross-correlation Analyses Confirm that Onset of Rainfall Triggers Mating Flights, not Peak Rainfall

Both the expert and literature surveys (Tables S3 and S4) predict that *Atta* mating flights are triggered by the first major rains at the beginning of a rainy season, not by peak rainfall of a rainy season. We tested this prediction by quantifying the time-lag between mating-flight peak and rainfall peak in cross-correlation analyses comparing time-series between rainfall and mating-flights in Colombia. By-month cross-correlation analysis revealed that rainfall and mating flights peak in the same month, with the highest correlation observed at lag = 0 months, indicating that the seasonal peaks of both variables are temporally congruent at the monthly scale (Figure S3A). The overall monthly correlation between rainfall and mating-flight activity is r=0.42±0.4 (mean±SD), indicating a moderately positive correlation. Six ecoregions had sufficient alate observations to permit calculation of within-ecoregion Pearson correlations between rainfall and mating-flight activity. Four of these six ecoregions (Dry Forests, Llanos, Moist Forests, Montane Forests) showed positive correlations (r = 0.45–0.79), while two ecoregions showed correlations near zero (Xeric Scrub r=−0.05; Caribbean Mangroves r=−0.0003). A one-sample t-test on these six correlations (testing against the null expectation of a value of r=0) indicated that rainfall and mating-flight activity were significantly positively correlated on average across these six ecoregions (t=2.82, df=5, p=0.019).

The mean Pearson correlation between daily rainfall and mating-flight activity across ecoregions was r=0.14±0.14 (mean±SD), suggesting a weak positive correlation. For the six ecoregions with sufficient alate observations (see above), a one-sample t-test on the six within-ecoregion Pearson correlations indicated that their overall average was significantly greater than zero (t=2.39, df=5, p=0.031), likewise suggesting, as in the above by-month analysis, a positive association between rainfall and mating-flight activity on average across ecoregions. The overall correlation is stronger in the cross-correlation analysis by month (r=0.42; see above) than by day (r=0.1), likely because there are many days during the year with zero flight observations even during months with peak mating activity, but that does not mean that *Atta* alates never fly on those specific days, but that no alate observations were recorded.

The daily cross-correlation analysis revealed that the lag of peak correlation between rainfall and mating-flight activity varied considerably across ecoregions, ranging from -5 days (Dry Forests) to -28 days (Llanos, Moist Forests, Xeric Scrub). However, the longer lag estimates of several weeks are less reliable because of the few flight observations available and the weak correlations estimated for those ecoregions (r < 0.17). In ecoregions with strongest signals (Dry Forests r=0.25; Montane Forests r=0.38, Figure S3B), mating flights peaked 5-10 days before corresponding rainfall peaks, confirming the above prediction that the onset of a rainy season triggers mating flights, rather than peak rainfall of a rainy season. While many observers noted that it is the first sufficient rainfall in a rainy season that triggers *Atta* mating flights (Tables S3&S4), the cross-correlation analyses here are the first statistical tests comparing time-series between rainfall and *Atta* mating-flights.

### 3.3 North-East and North-Central South America

Other interesting biogeographic regions across the *Atta* range are north-central South America and north-eastern South America (i.e., north-eastern Brazil). In our maps averaging rainfall across 2012-2021 (Figure 2), north-east Brazil is noticeably drier, consistent with known semi-arid conditions in north-east Brazil (Alvares et al. 2013) and with known drought adaptations of *Atta opaciceps*, the dominant *Atta* species in seasonally arid regions of north-east Brazil (Gonçalves 1951; Schaefer at al. 2021). Marínho et al. (2011, page 167) write that the “hottest part of Brazil is the northeast, where temperatures of more than 38 °C (100 °F) are frequently recorded during the dry season between May and November.” This corresponds to a period from June to September with near-absence of observations at iNaturalist of *Atta* reproductives in north-east South America (Figure 1G), followed by an extended flight season spanning half a year from November to April (Figure 1G). The 110 records of *Atta* reproductives in the Insect Collection of the Cocoa Research Center (CEPLAC) in Bahia, Brazil (Questionnaire #30, Table S3) show (i) a clear March-May peak in *Atta* mating-flights in north-east Brazil, (ii) a seasonal low (but not complete absence) of *Atta* mating-flights in July-October, (iii) a possible increase in *Atta* mating-flights in November, and therefore (iv) likewise an extended mating-flight season of at least 3-6 months (SI-1 Footnote 3). Unlike the iNaturalist dataset where most observations cannot be identified to species, the CEPLAC dataset includes only records of identified *Atta* species, allowing species comparisons. For north-east Brazil, there is no clear indication that the four *Atta* species represented in the CEPLAC dataset have different annual mating-flight phenologies (Table S5), but sample sizes are small for *A. opaciceps* (n=7) and *A. laevigata* (n=5), and the only robust comparison between *A. cephalotes* (n=67) and *A. sexdens* (n=31) suggests comparable mating-flight phenologies in north-east Brazil for these two *Atta* species.

For north-central South America (Figures 1E&F), the sample sizes of iNaturalist observations of *Atta* reproductives are very small (n=17 in total), with most of these from the State of Amazonas in Brazil from the Manaus area (n=11, October-December), some from the State of Pará (n=5, January-May), and only one observation from the Guianas (n=1, December, French Guyana). These represent too few observations to infer patterns of *Atta* mating flights in this region, except perhaps for Manaus where flights in our dataset were restricted to October-December with a peak in November (SI-1 Footnote 4).

### 3.4 Transect Colombia–Central America–México–USA

Throughout Central America, México and the USA, *Atta* have unimodal mating-flight phenology (Figures 1A-C). After a drier period in winter in this region, the onset of *Atta* mating flights (Figures 5&6) each year correlates with the geographically-varying onset of rains. Because the onset of rains moves across this area from south-east to north-west (Figure 5 inserts), the peak of *Atta* mating flights is delayed by two months in north-westerly México (July) relative to south-easterly Central America (May; compare Figures 5B-5D with Figures 5L-5K). Specifically, the large sample size of observations across Central and North America (nearly 1200 observations; Table S1) enables us to elucidate mating-flight phenology on a more fine-grained scale (Figure 5). The shifting rains generate systematic, concomitant variation of *Atta* mating-flight phenology along a 4800-kilometer transect, ranging from north-west Colombia (Figure 5A) to Arizona, USA (Figure 5K). Rains trigger mating flights earliest in Colombia (April) and Central America (May), latest in north-west México (July) and Arizona, USA (July, August). A parallel correspondence between rainfall and *Atta* mating flights exists also along a shorter transect along the Gulf Coast from the Yucatan (Figure 5L) to southmost USA (Figure 5O).

**FIGURE 5.**
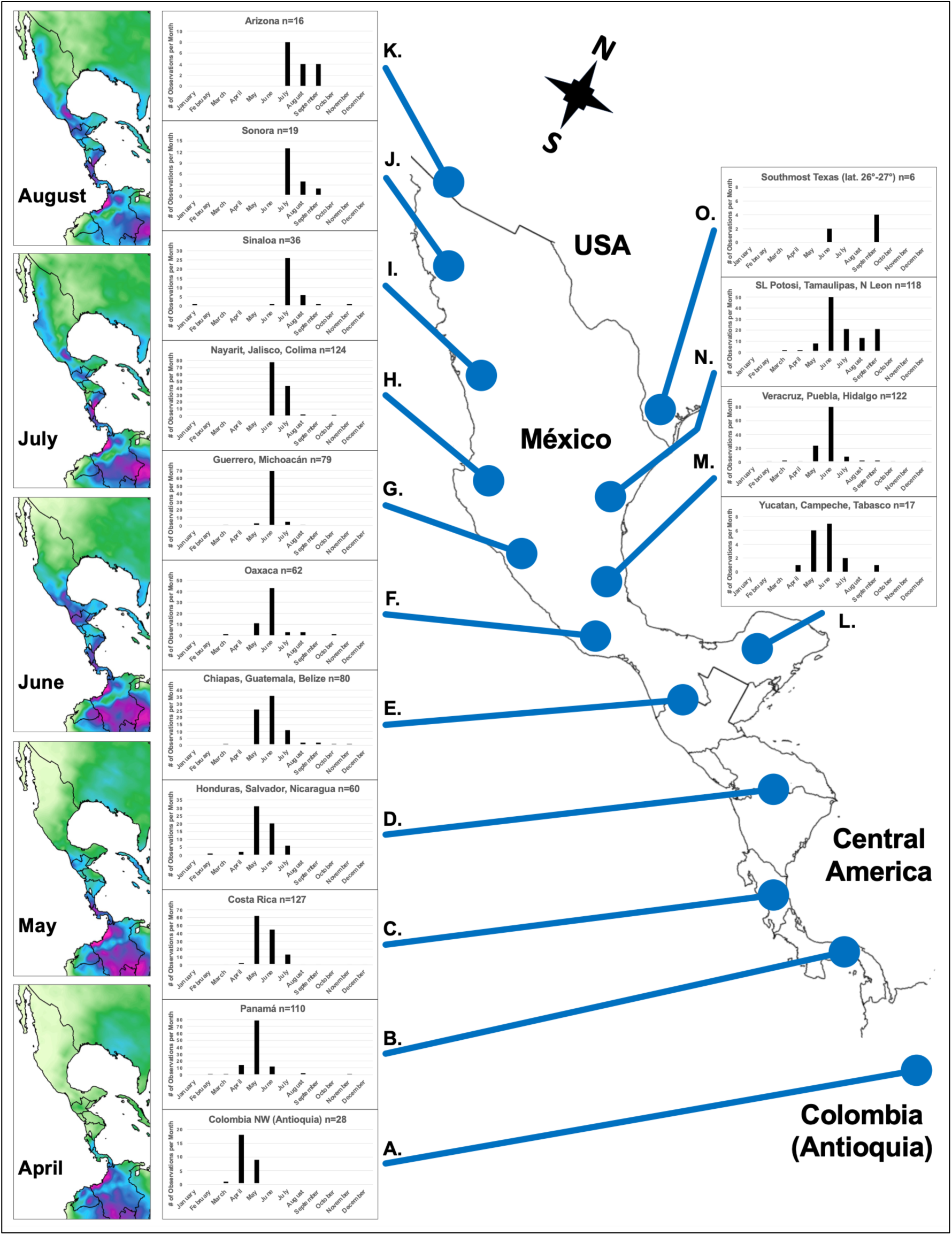
Mating-flight phenology of *Atta* leafcutter ants (n=1191 observations) along a 4800-kilometer transect from north-west Colombia (**A**) across Central America (**B**-**E**) and México (**E**-**J**) to Arizona, USA (**K**). Mating-flight phenology changes systematically along this transect, corresponding to the gradual spread of rains during spring and summer along this transect (see the rainfall timeline illustrated in inserts at left; these maps are cropped from Figure 2). Rains trigger *Atta* mating flights earliest in Colombia (April) and Central America (May), latest in north-west México (July) and Arizona (July, August). A parallel correspondence between rainfall and *Atta* mating flights exists also along a shorter transect along the Gulf Coast from the Yucatan (**L**) to southmost USA (**O**). The predominance of *Atta* mating flights in May in Central America (**B**-**D**) explains why *Atta* reproductives are called “hormigas de Mayo” (May ants) throughout Central America (SI-1 Footnote 5).

To further explore *Atta* mating-flight phenology along the transect from Central America to north-west México (Figure 5), we calculated for each observation the ordinal date (continuous count of days in a year, starting on 1^st^ of January as Day 1), then estimated the trend between ordinal date and latitude of observations (Figure 6A), and between ordinal date and longitude of observations (Figure 6B). We estimated the onset of mating flights within a given latitudinal bracket (or alternatively longitudinal bracket) using only the 30% of the earliest observations (black dots in Figures 6A&B, corresponding black trendlines) within a given bracket. The trendlines in Figures 6A&B confirm the pattern visualized also in Figures 5A-5K that, along the south-east to north-west transect, the onset of *Atta* mating flights is earliest in Panamá and Costa Rica (May), and latest in north-west México (July), corresponding to the gradual spread of spring rains from south-east to north-west (see rainfall map inserts in Figure 5) along this 4800-kilometer transect.

**FIGURE 6.**
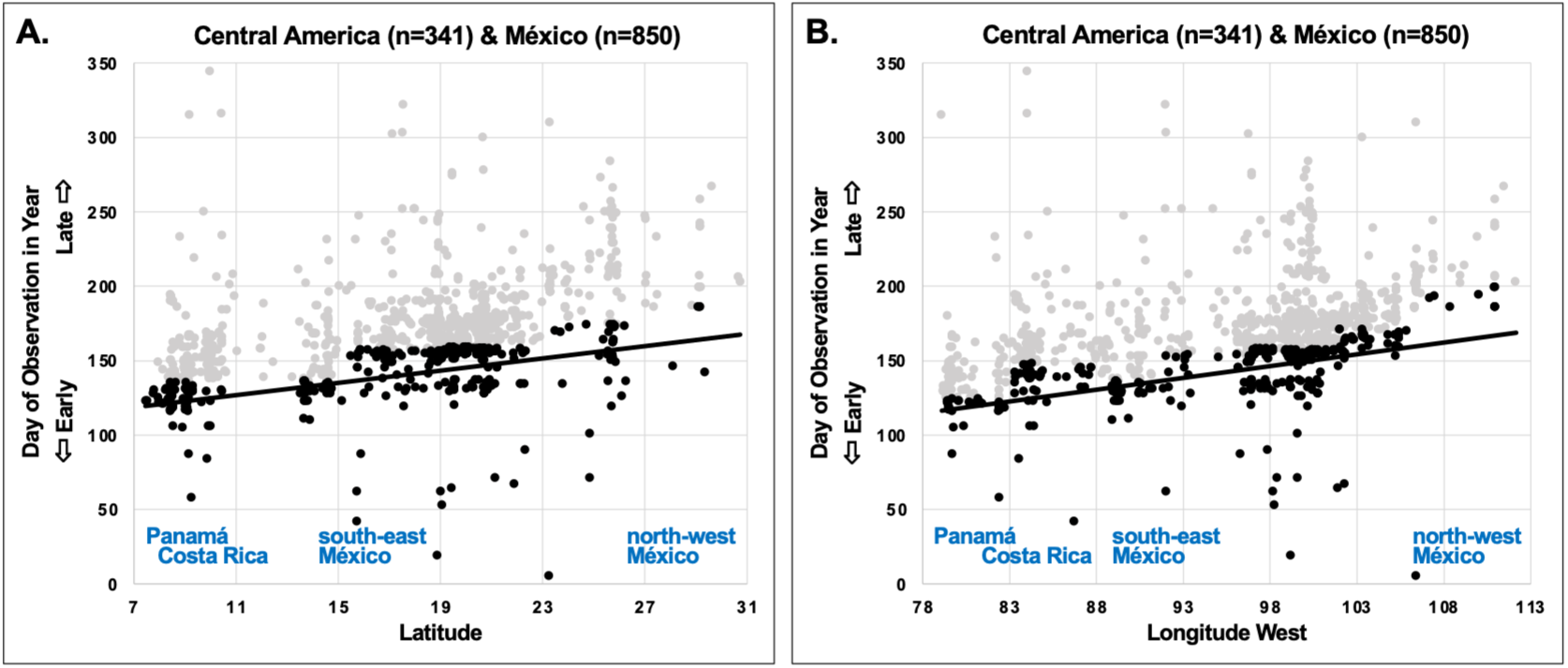
Latitudinal (**A**) and longitudinal (**B**) gradients of *Atta* mating-flight phenology across Central America and México, estimated from observations of *Atta* reproductives in 2012-2021 at iNaturalist. The ordinal date (numerical day in year) of 1191 observations (n=341 observations from Central America; n=850 from México) is plotted against latitude or longitude. **A. Latitude:** 30% of the earliest mating-flight records in spring (black circles, black trendline; Spearman rho=0.615, p<0.00001) observed for *Atta* in any given latitudinal bracket (e.g., N7-11°, N11-15°, etc.), relative to the remaining 70% of later mating-flight records (gray circles) observed for each of the same latitudinal brackets. **B. Longitude:** 30% of the earliest mating-flight records in spring (black circles, black linear trendline; Spearman rho=0.738, p<0.00001) observed for *Atta* in any given longitudinal bracket (e.g., W78-83°, W83-88°, etc.), relative to the remaining 70% of later mating-flight records (gray circles) observed for each of the same longitudinal brackets. Longitude is plotted in absolute values as Longitude West. **A & B:** The trendlines estimate onset of mating flights in spring using an arbitrary 30% cutoff criterion for earliest records; analyses using other cutoff criteria (e.g., 20% or 50%) show the same trends (graphs not shown). For both latitudinal and longitudinal gradients, therefore, the spring onset of mating flights is earliest in Panamá and Costa Rica, and latest in north-west México, corresponding to the annual onset or rains spreading from south-east to north-west across that region (Figures 2 and 5). See SI-1 Footnote 6 for additional discussion.

### 3.5 Within-Species Mating-Flight Phenology of *Atta texana*

Because only *Atta texana* occurs in Louisiana and Texas, we were able to evaluate within-species mating-flight patterns for this single species, using here our extended dataset (n=592) that included also iNaturalist observations from 2022-2024. Whereas all other datasets discussed so far cover the 10-year timespan 2012-2021, the *A. texana* dataset here covers the 11-year period from 2014-2024 (we found no records at iNaturalist of *A. texana* reproductives from before 2014). Figure 7 plots the ordinal observation date in a year for all 592 observations against latitude (Figure 7A) and longitude (Figure 7B). Mating flights of *A. texana* occur at dawn following a day of sufficient rainfall (minimum rainfall of ≈1.0-1.5 cm is necessary to stimulate flights of *A. texana*; UGM personal observation), and the onset of *A. texana* mating flights tends to occur later in more western longitudes (Figure 7B) where spring rains with sufficient precipitation arrive later in spring (May/June) compared to typically April/May in central Texas. In addition to sufficient rainfall, mating flights of *A. texana* are also dependent on temperature at dawn on the morning following sufficient rainfall (minimum temperature at dawn of about 16 °C is necessary to permit mating flights in *A. texana*; Moser 1967; Questionnaire #35 in Table S3). Dawn temperatures increase gradually during the spring months of March-June across the southern USA, and the onset of mating flights occurs therefore later in the more northern populations of *A. texana* (Figure 7A) where dawn temperatures are sufficiently warm only later in spring. Hart et al. (2018) documented a similar latitudinal trend for mating flights of all ant species observed in 2012-2014 in a citizen-science project in the United Kingdom, where flights occurred later in the season at higher latitudes.

**FIGURE 7.**
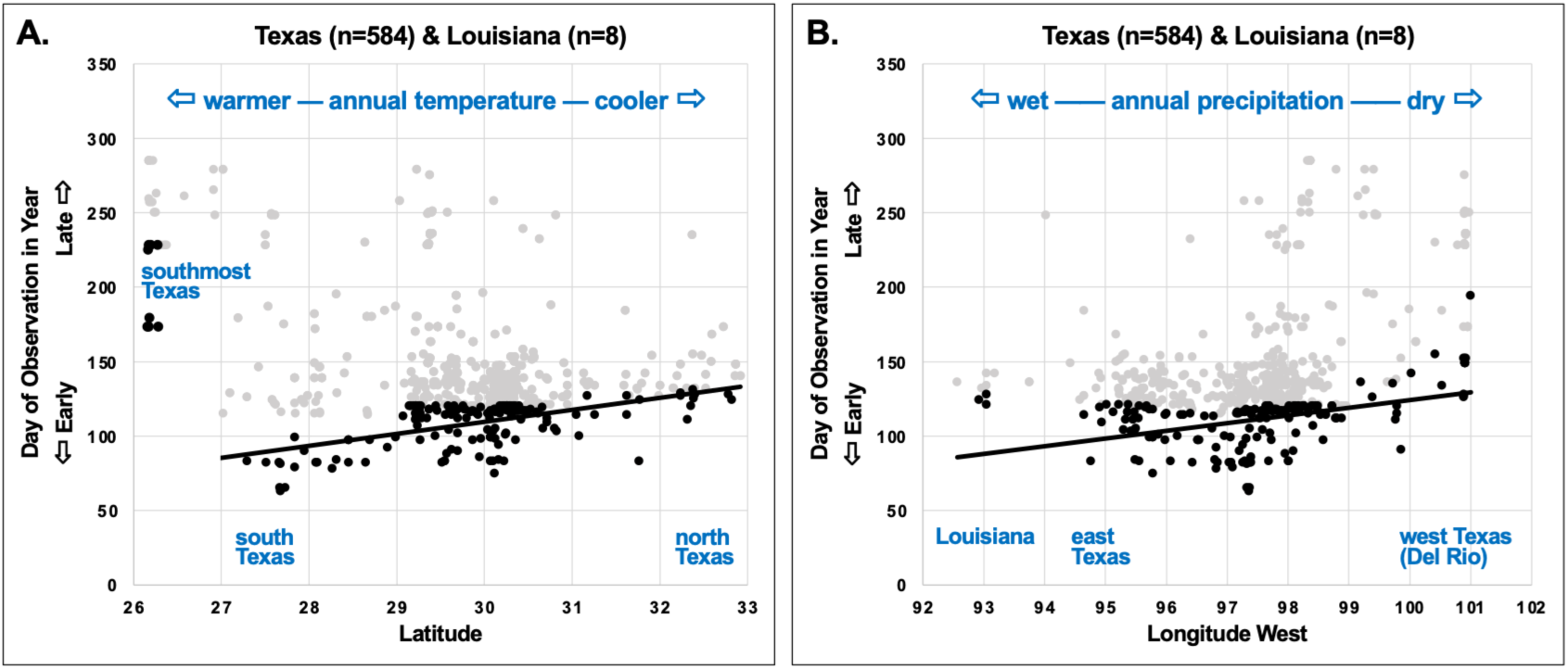
Latitudinal (**A**) and longitudinal (**B**) gradients of *Atta texana* mating-flight phenology across Texas and Louisiana, estimated from observations of *A. texana* reproductives in 2014-2024. The ordinal date (numerical day in year) of 592 observations (n=584 observations from Texas; n=8 from Louisiana) is plotted against latitude or longitude. **A. Latitude:** 30% of the earliest mating-flight records in spring (black circles, black linear trendline) observed for *A. texana* in any given latitudinal bracket (e.g., N26-27°; N27-28°; etc.), relative to the remaining 70% of later mating-flight records (gray circles) observed for each of the same latitudinal brackets. *Atta* reproductives in southernmost Texas between latitudes N26-27° (south-eastern “lower” Rio Grande Valley) have so far not been observed in spring, only later between late June and October (see Figure 8 for further analysis of these late mating flights in southmost Texas). For any latitudinal bracket north of latitude N27°, in contrast, the earliest mating flights occur in spring, but increasingly later at higher latitudes, corresponding to the gradual warming during spring of dawn temperatures when *A. texana* has its mating flights. For the range between latitude N27-33° (i.e., excluding the very late-flying populations at latitude N26-27° in south Texas), the correlation between ordinal date and latitude is statistically highly significant (Spearman rho = 0.429, p<0.00001). **B. Longitude:** 30% of the earliest mating-flight records in spring (black circles, black linear trendline) observed for *A. texana* in any given latitudinal bracket (e.g., W92-93°; W93-94° longitude, etc.), relative to the remaining 70% of later mating-flight records (gray circles) observed for each of the same longitudinal brackets. The correlation between ordinal date and longitude is statistically highly significant (Spearman rho = 0.431, p<0.00001). Mating flights occur somewhat later towards the western range limit of the *A. texana* distribution (Del Rio area), corresponding to the later annual arrival of rains in late spring and early summer in west Texas. **A & B:** The trendlines estimate onset of mating flights in spring using an arbitrary 30% cutoff criterion for earliest records; analyses using other cutoff criteria (e.g., 20% or 50%) show the same trends (graphs not shown).

Surprisingly, *A. texana* reproductives in southernmost Texas between latitudes N26-27° (south-eastern “lower” Rio Grande Valley) have so far not been reported at iNaturalist in spring, only later in the year between late June and October, whereas at higher latitudes between N27-33°, *A. texana* mating flights were observed primarily in March-June (Figures 7A&8; SI-1 Footnote 7). This finding prompted an analysis of mating-flight phenology along a south-to-north transect from Mexico to Texas (see 3.6).

**FIGURE 8.**
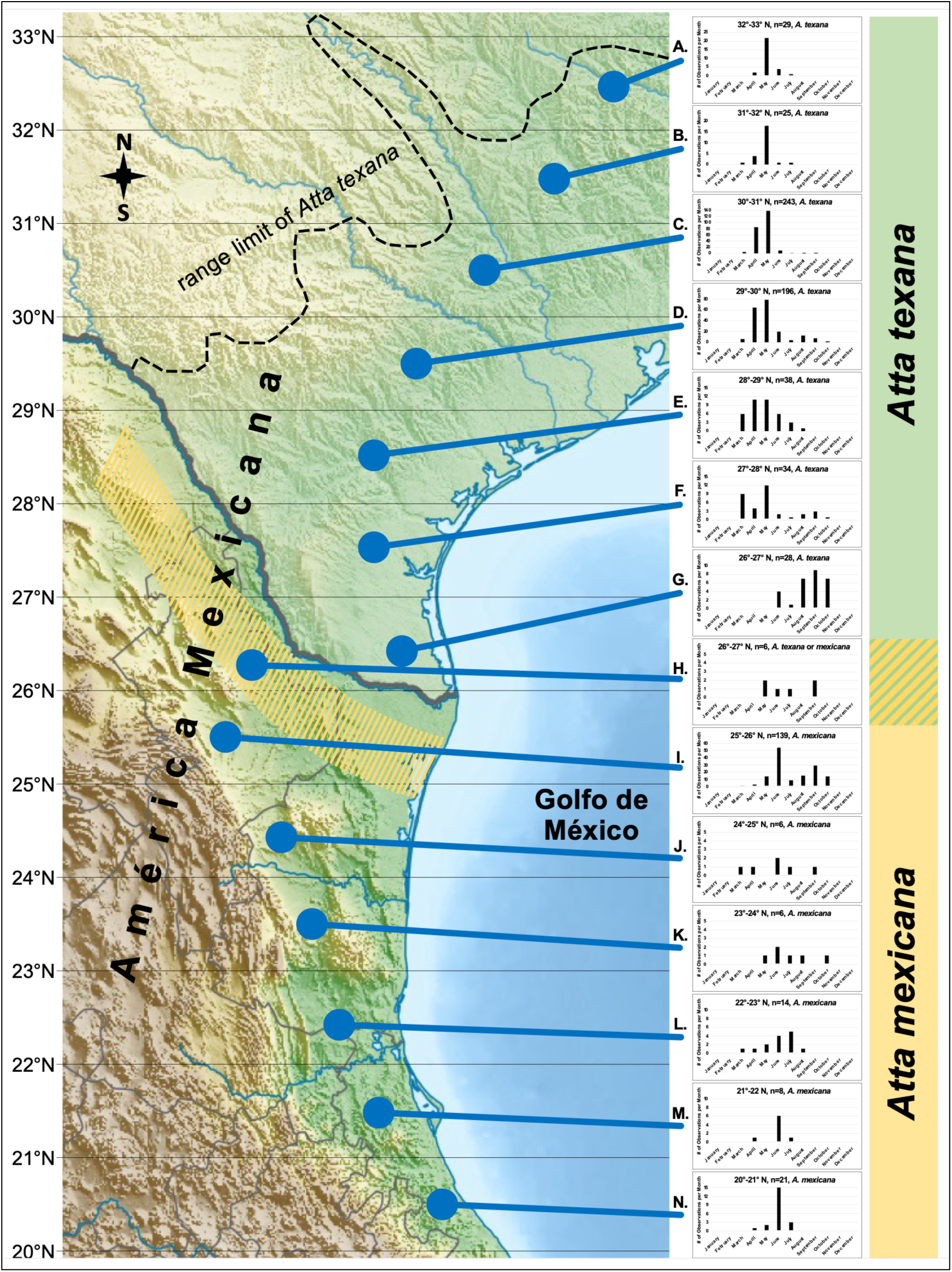
Mating-flight phenology of *Atta mexicana* and *A. texana* leafcutter ants along a 1500-kilometer transect from the State of Veracruz in México northward along the Gulf coast to the northernmost *Atta* populations in northern Texas, USA. Phenology graphs **A.-N.** include all records of *Atta* reproductives from 2012-2024 available at iNaturalist. Only *A. texana* occurs in Texas, and no *A. mexicana* have so far been found in southmost Texas (**G.**). Only *A. mexicana* appears to occur along the Gulf Coast in México, ranging south from the Monterrey area (**I.**) to the northern half of the State of Veracruz (**N.**). *A. texana* and *A. mexicana* are likely sympatric in north-east México in north-east Coahuila, northern Nuevo León (**H.**) and northern Tamaulipas (no records of *Atta* reproductives are available at iNaturalist for that biogeographically interesting region in Tamaulipas). The most likely area of sympatry between *A. mexicana* and *A. texana* is indicated in the map as a yellow-striped band somewhat south of the Rio Grande River. Only records from the Gulf-Coast lowlands in México are included in the phenology graphs here (i.e., records from the more mountainous areas to the west of the Gulf Coast lowlands in México are not included here), because *Atta* mating flights occur slightly later in the year in the more western mountainous areas than in the Gulf Coast lowlands (data not shown). Records from México south of latitude N20° (southern half of the State of Veracruz and further south) are not included in the transect because south of latitude N20° *A. mexicana* is sympatric with *A. cephalotes*. Between latitudes N27° and N33° in Texas, *A. texana* has mating flights primarily in spring, with the more southern populations somewhat earlier in March-May (**F. & E.**) and the northern populations later in May-June (**A. & B.**). Between latitudes N20° and N26° in México, *A. mexicana* has mating flights in summer and fall, with the more southern populations somewhat earlier in June-July (**L. & N.**) and the northern populations later in June-October (**I. & K.**). Interestingly, in southmost Texas, mating flights of *A. texana* have been observed so far only in June-October (**G.**, n=28 records at iNaturalist, this includes one *A. texana* record from Reynosa at the México side of the Rio Grande River), corresponding in southmost Texas more to the summer-fall mating-flight phenology of *A. mexicana* (**I.-N.)** than to the spring mating-flight phenology (**A.-F.**) typical for the rest of the *A. texana* range. Relief map from Wikimedia, CC BY-SA 3.0.

### 3.6 Transect Across Contact Zone Between *Atta mexicana* and *Atta texana*

Figure 8 shows latitudinal variation in mating-flight phenology across a contact zone between *Atta mexicana* and *A. texana* in far north-east México and the neighboring Texas, USA. Throughout north-east México, *A. mexicana* has mating flights later in the year, primarily in June and July (Figures 8J-8N), whereas somewhat further north in southern Texas (Figures 8D-8F), *A. texana* has mating flights in spring between March-May, except for southmost Texas (Figure 8G) where *A. texana* mating flights have been observed only later in the year, just as for *A. mexicana* somewhat further south. The exact area of sympatry between *A. mexicana* and *A. texana* is insufficiently known (it is not in Texas because *A. mexicana* does not occur there), but *A. mexicana* and *A. texana* are likely sympatric somewhat south of the Mexico-USA border (see band of likely sympatry in Figure 8; SI-1 Footnote 8 summarizes all relevant evidence).

Interestingly, the transition in Figure 8 from primarily late-flying *A. mexicana* (summer) to primarily early-flying *A. texana* (spring) occurs only approximately in the region of most likely sympatry, but does not coincide with the area of sympatry as expected if mating-flight season evolved by character displacement. Both species have mating flights at dawn following sufficient rainfall, reproductives of the two species therefore could potentially interact at dawn in areas of sympatry, and the specialization on spring mating by *A. texana* versus summer mating by *A. mexicana* could therefore be interpreted as a possible prezygotic isolation mechanism. However, going south to north along the transect in Figure 8, the actual transition from mating flights later in the year to mating flights in early spring occurs actually *within* the species range of *A. texana* (compare Figures 8F&G), and somewhat north of the contact zone between *A. mexicana* and *A. texana*. Prezygotic reproductive isolation and possible character displacement therefore does not seem to explain the mating-flight transition within *A. texana*, and other factors — perhaps related to a combination of precipitation, temperature, and nutrients available in spring vegetation cut by *Atta* — could explain the transition within *A. texana* in southmost Texas (see additional discussion in section 5.4 on character displacement). Because different populations of *A. texana* in southmost Texas have mating flights later in the year than populations in the rest of the *A. texana* range, it is possible that the southmost *A. texana* populations are genetically more distinct than predicted by isolation-by-distance.

## 4 Key Results

*Atta* leafcutter ants are bioindicators with mating phenologies that respond at fine scales to geographic and seasonal climate variation across the Americas. This new *Atta* bioindicator system offers several advantages. First, because of the continuous distribution of *Atta* across the Americas from S33.6° to N33.2° and covering diverse habitats except higher elevations, *Atta* mating-flight phenologies track climate comprehensively across a 9200-kilometer trans-tropical transect spanning both the Southern and Northen Hemispheres (Figures 1&2). Second, because onset of mating flights can be timed with great precision in *Atta* populations when synchronized mass-mating flights are triggered by the first major rainfall of rainy seasons, *Atta* bioindicators are (1) accurate sensors of precipitation changes across the tropics and subtropics; and (2) accurate sensors of both temperature and precipitation changes at their range limits in the southern USA (Figures 7&8) and in Uruguay/Argentina (Kusnezov 1962). Third, the community database iNaturalist accumulates increasingly more (Table S2) detailed records of *Atta* mating-flight phenologies across the entire *Atta* range, and we verify here the accuracy of these mating-flight patterns inferred from iNaturalist data by comparison with (i) mating-flight data reported in the *Atta* literature, and (ii) mating-flight data accumulated by a consortium of experts who have researched *Atta* for a combined 1000+ work-years (Figure 1). We illustrate the climate sensitivity of *Atta* bioindicators in case studies of latitudinal and longitudinal climate transects across Central America and Mexico (Figures 5&6); latitudinal and longitudinal climate transects across the southern USA (Figures 7&8); and regional ecosystem variation within Colombia (Figures 3&S2), where regional differences in *Atta* mating-flight phenology correspond to temporal differences in rainfall between ecoregions of Colombia (Figure 4).

## 5 Conclusions and Future Research Directions

Climate-dependent mating-flight phenology in *Atta* is a valid metric to elucidate spatial and temporal climate changes across the Americas. *Atta* mating-flight phenology can therefore serve as an indicator to (1) evaluate climate impacts on many other arthropod species whose biologies are likewise tied to precipitation and temperature, and (2) quantify climate impacts on life-history traits expected under future global climate change. A key advantage of the *Atta* bioindicator system is the continuously growing stream of iNaturalist records. Specifically, for our analysis of *Atta* mating phenologies across the Americas, we explore here only iNaturalist observations from 2012-2021 (n=2335; Figure 1), but we estimate that subsequently, for the years 2022-2026, at least 2.5-times additional observations (Table S2) of *Atta* mating flights were submitted to iNaturalist. This suggests that climate-dependent mating-flight phenology of *Atta* can be studied in the future at much finer geographic and temporal scales than we are able to resolve here with our 2012-2021 dataset, particularly for the biogeographically most interesting regions of Colombia/Ecuador, north-east Brazil, the Northern-Hemisphere transect from Colombia to the USA (Figure 5), and the southern and northern range limits of *Atta* where mating-flight phenology is dependent on both temperature and precipitation (Figure 7). We suggest the following specific projects for future *Atta* research linking climate, biogeography, and life history:

### 5.1 Unexplored Climate Gradients

Future research should use the continuously growing iNaturalist information on *Atta* to address so far unexplored gradients in precipitation, temperature, and altitude, such as (i) the east-west precipitation gradient in Uruguay/Argentina, where spring rains start earlier in the east and somewhat later in the west (Figure 2) and where the east-west precipitation gradient is oriented, as in the south-central USA, orthogonal to a south-north temperature gradient, such that the independent impacts of temperature and precipitation changes on mating-flight phenology can be partitioned in statistical analyses; (ii) a south-to-north precipitation gradient from Uruguay across Brazil to the Guianas, where spring rains start earlier in the south than in the north (Figure 2); and (iii) elevational gradients, for example along the Andean slopes in Colombia, Ecuador, and Peru; or elevational transects from the Atlantic to the Pacific lowlands across the central mountain ranges in Panamá, Costa Rica, Nicaragua, Guatemala, and México.

### 5.2 Cyclical Climate Change and Future Climate Trends

Future research will be able to expand on our *Atta* datasets (Data S1-S4; Tables S3-S5) to quantify phenology responses to climate cycles and to multi-decade climate trends. Specifically, because the range of *Atta* covers tropical, subtropical, and some temperate climate zones from S33.6° to N33.2° in the Americas, *Atta* bioindicators should be useful to track impacts of cyclical climate changes driven by the El Niño-Southern Oscillations and the Atlantic Intertropical Convergence Zone Oscillations. Such research will benefit from the many entomology collections throughout the Americas that curate records of *Atta* reproductives, as well as from detailed *Atta* mating-flight records that predate iNaturalist, to establish ant-phenology observatories sensu Kusnezov (1962, page 441), for example at the Reserva Ecológica El Bagual in Argentina (Questionnaire #01, Table S3); Botucatu, Rio Claro, and Viçosa in southern Brazil (Questionnaires #05, #13, #18, #26, #34), the Cocoa Research Center in north-east Brazil (Questionnaire #30), Colombia (Questionnaires #07, #15), and south-central USA (Questionnaires #08, #35).

### 5.3 Predictable shifts in mating-flight timing and synchrony

The latitudinal synchronicity hypothesis (Kaspari et al. 2001) predicts for ants that mating seasons should be shorter and more seasonal at higher latitudes, because the prolonged winters there delay mating flights in spring (Kaspari et al. 2001; Dunn et al. 2007; Helms 2023). Our analyses support that prediction for higher latitudes in the south-central USA (Figures 7A&8), but not for higher latitudes in México (Figures 5&6A). If climate continues to warm in the future, mating flights are predicted to advance in spring and possibly become less synchronized in the currently northernmost populations of these two *Atta* species. Quantitatively, spring advancement predicts more shallow slopes (reduced steepness) of the trendlines for spring onset of mating-flight shown in Figures 6A&7A, because spring advancement under climate warming should be greater at higher latitudes than at lower latitudes. Future analyses will be better able to test these predictions when including iNaturalist observations from additional years not included in our 10-year dataset. Parallel analyses of the *Atta* populations across Uruguay/Argentina, Paraguay, and southern Brazil will permit a separate test of the latitudinal synchronicity hypothesis for the southern range limit of *Atta*, which we did not test here because of the paucity of current iNaturalist observations from these southernmost regions. We also did not explore altitudinal trends (delayed flight seasons are predicted for higher altitudes), but sufficient iNaturalist observations may already exit in our dataset (Data S1) to evaluate transects from the Atlantic to the Pacific lowlands across the central mountain ranges in Central America and across México. Lastly, whereas mating-flight phenology appears less seasonal and more prolonged in equatorial latitudes of the *Atta* range in Figure 1 compared to the shorter mating-flight seasons at higher latitudes (compare aseasonality in Figures 1D-1G with seasonality in Figures 1A&1H), only a rigorous test using circular statistics of mating-flight seasonality (Tozetto et al. 2023) will be able to quantify such latitudinal differences in seasonality.

### 5.4 Sympatry and Character Displacement in Mating-Flight Phenology

In leafcutter ants, a major shift in mating-flight season has been found so far only in a social-parasitic species (*Acromyrmex charruanus*), which has been interpreted as a possible pre-zygotic reproductive barrier between the social parasite and its closely-related host (Rabeling et al. 2015) and a possible case of character-displacement evolution. To test for additional cases of character displacement using our *Atta* mating-flight datasets, the area of sympatry between *A. mexicana* and *texana* in north-east Mexico (Figure 8; Footnote 8) is the only location where we can evaluate possible character displacement in mating-flight phenology. *A. mexicana* and *texana* have mating flights shortly before dawn throughout their ranges, and there is no evidence that sympatric populations of the two species have mating flights in different months (Figures 8G-8I; Table S5). The two species presumably have mating flights on the same days at the same time-of-day in their area of sympatry, similar to other sympatric *Atta* species overlapping in mating-flight phenology in many other locations (Tables S3&S5). Differences in mating pheromones between *A. mexicana* and *texana* in the area of sympatry may be a possible pre-zygotic barrier preventing interbreeding between these two species.

### 5.5 Biogeography of Zeitgeber Cues Driving Production of Sexual Brood

To understand *Atta* mating-flight phenology on a behavioral-physiological level, it is insufficient to analyze only the environmental variables that trigger flights (sufficient rainfall) or that permit flights (minimum temperature) that we discuss above. Instead, because *Atta* colonies do not rear sexual individuals (alates) throughout the year, understanding the underlying behavioral-physiological mechanisms requires analysis of the Zeitgeber cues that cause *Atta* colonies to start rearing alate brood 2-3 months before the onset of flight-triggering rains, typically during drier and cooler months preceding wetter months with mating flights. It is nutritionally costly for an *Atta* colony to support thousands of alates in the nest, so colonies optimize energy budgets by initiating alate production 2-3 months before mating flights (i.e., the time needed by alates to complete development from egg to adult). To begin alate production months in advance of mating-flight-triggering rains, ant colonies are thought to rely on extrinsic Zeitgeber cues, which have not been investigated in *Atta*. Assuming that alate brood production follows an endogenous annual rhythm, as in other ant species (Kipyatkov 1995), potential Zeitgeber cues at higher latitudes could include seasonal changes in temperature or changes in daylength (the most reliable Zeitgeber synchronizing annual rhythmicity in animals). In contrast, at latitudes near the equator where daylength and temperature vary little throughout the year, potential Zeitgeber may be the climate transition from wet to dry season, or perhaps seasonally-varying nutritional cues of plants cut by workers. One of the most intriguing questions is, however, how Zeitgeber cues are perceived by the queens (e.g., to begin laying unfertilized eggs for production of male alates), because queens of mature *Atta* colonies do not leave the underground chambers so they cannot perceive daylength directly. Queens may respond directly to thermal changes experienced underground, or indirectly to Zeitgeber cues mediated by workers, by the cultivated fungus, or by both. Interactions between queen, workers, and fungus – modulated by photoperiod, thermoperiodicity, or food quality – are likely the primary behavioral-physiological mechanisms underlying differences in the timing of mating flights between regions and along biogeographic gradients documented in Figures 3-9.

### 5.6 Machine Learning of *Atta* Identification

Because most *Atta* records at iNaturalist do not specify whether the observation is an ant worker, a reproductive, or a nest, we had to inspect all *Atta* images individually, then copy respective information on *Atta* mating flights into our custom spreadsheet (Data S1). A more expedient future approach will be to use machine learning to automate this entire process, for example by using a portion of our well-curated dataset (Data S1) to train an image-recognition algorithm, test this trained algorithm on the remaining portion of our dataset, use the recursively-trained image-recognition algorithm to assemble a continuously growing iNaturalist dataset of *Atta* reproductives, then validate by visual inspection only those records that are identified by the trained algorithm. We estimate that the average growth rate of new observations of *Atta* reproductives at iNaturalist was about 15%/year in the years 2021-2025 (Table S2), that there are currently nearly 6,000 records of *Atta* reproductives deposited at iNaturalist (more than twice the dataset of n=2335 analyzed in our study here), and that there will be more than 17,000 records overall of *Atta* reproductives at iNaturalist by 2030 (assuming future contributions to iNaturalist at current growth rates; Table S2). To update our dataset (Data S1) by adding future observations, it will be increasingly more expedient to automate the process of image recognition and data extraction.

### 5.7 Life-History Information Archived by Ethnography and by Entomology Collections

In our survey of publications on *Atta* mating-flight phenology, we found many detailed descriptions of *Atta* mating flights in the ethnozoological literature (Table S4). Historically, ethnographers were the first to describe *Atta* mating flights nearly 500 years ago (e.g., Anchieta 1560), because *Atta* female reproductives are collected for food by indigenous peoples (Dufour 1987; Costa Neto and Ramos-Elorduy 2006; Aguilera-Espinosa et al. 2024). Centuries of ethnography of Amerindian food traditions documented remarkably detailed knowledge accumulated by indigenous peoples of *Atta* mating-flight biology, including indigenous knowledge of species differences (e.g., sympatric leafcutter species are distinguished by unique names in many Amerindian languages), habitat preferences of different *Atta* species, their species-specific diurnal time windows when each species has its mating flights, and the specific climate conditions triggering mating flights (Table S4). Likewise, entomology collections preserving insect specimens are repositories of a wealth of biological knowledge, including information on mating-flight phenology (see Questionnaires #07 and #30; Kusnezov 1962). We therefore predict that integrated efforts involving ethnographers, museum curators, and entomologists will further optimize *Atta* as high-resolution bioindicators of climate across the Americas.

## Author Contributions

U.G.M., D.M.R., T.D.K., and F.R. conceived the study. U.G.M., T.D.K., S.R., R.V.B., M.N.B., R.E., and K.I.-L. curated data from iNaturalist. U.G.M., D.M.R., F.C.S.R., and T.G.M. performed analyses and produced figures. T.G.M., D.M.R., and F.C.S.R. prepared climate data for climate figures. F.R. and U.G.M. developed the Expert Survey. U.G.M. performed the literature review with support from many co-authors.

U.G.M. wrote the first draft of the manuscript. F.R., D.M.R., F.C.S.R., T.G.M., C.R., J.M.L., R.I.S., J.S.C., H.L.V., A.R., M.B.Jr, and H.F.-M. edited and expanded the manuscript. All *Atta*-expert co-authors contributed observations to the Expert Survey, evaluated the validity of the iNaturalist data, and commented on the manuscript.

## Acknowledgements

We thank the thousands of citizen scientists who contributed observations through iNaturalist; K. Rao for help with databasing information from iNaturalist; C. Aliaga, S. Arcuri, R. Ochoa, F. Pagnocca, and J. Ticona for sharing unpublished records of *Atta* mating flights; and anonymous reviewers for many helpful suggestions to improve the manuscript.

## Data and Code Availability Statement

iNaturalist data are publicly available (inaturalist.org). The Supporting Information includes all analyzed datasets in Data S1-Data S5 and all code in SI-3 R-scripts.

**Additional information is online in the Supporting Information (SI) files:**

**SI-1:** Expanded methods, footnotes, and supplementary references.

**SI-2:** GIF of monthly precipitation maps.

**SI-3:** R-scripts S1-S4.

**Figure_S1:** Maps of average monthly precipitation in 2012-2021 across the *Atta* range.

**Figure_S2:** Atta mating-flight phenologies in Colombia inferred from iNaturalist & UNAB data.

**Figure_S3:** Cross-correlations between rainfall and *Atta* mating-flight activity in Colombian ecoregions.

**Table_S1:** Number of iNaturalist observations, by country and by *Atta* species, in Data_S1.

**Table_S2:** Annual number of iNaturalist observations in Data_S1.

**Table_S3:** Questionnaire survey of expert observations of *Atta* mating-flight biology.

**Table_S4:** Literature survey of *Atta* mating-flight biology.

**Table_S5:** Combined data from Tables S3 & S4, grouped by region and by identified *Atta* species.

**Data_S1:** iNaturalist records of *Atta* reproductives.

**Data_S2:** Data compiled from Table S3 expert survey.

**Data_S3:** Data compiled from Table S4 literature survey.

**Data_S4:** Colombia data combined from UNAB records and iNaturalist records.

**Data_S5:** Colombia precipitation data from Urrea et al. 2019.

