## Supporting Information for "*Atta* Leafcutter Ants are Fine-Scale Bioindicators of Geographic and Seasonal Climate Changes Across the Americas"

**INDEX**

Pages 2-5 Methods, Additional Details

Pages 6-8 Footnotes 1-7

Pages 9 References cited in Supporting Information

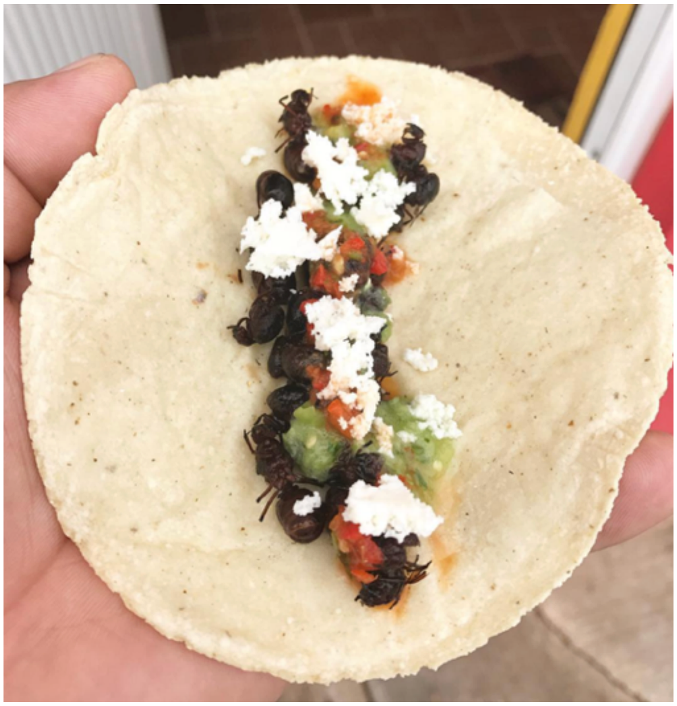

### METHODS, ADDITIONAL DETAILS

#### Data collection

During the first two years of the Covid pandemic (2020-2021), when we had to conduct research remotely because of Covid guidelines, undergraduate research assistants at the University of Texas at Austin evaluated observations of *Atta* reproductives (alate females, dealate females, or alate males) deposited at iNaturalist (inaturalist.org), then import these observations into our customized Excel spreadsheet (Supporting Information Data S1). Research assistants were instructed to filter observations at iNaturalist using the Filter Option set on a specific day by *Date Added* (date on which an observation had been submitted to iNaturalist by an observer) and sort by ascending order of the date of submission (i.e., essentially, sort by serial number at iNaturalist), starting with the first *Atta* reproductive record in February 2012 (https://www.inaturalist.org/observations/54085). If a record included multiple images that had been combined by an observer into a single observation event, assistants were asked to evaluate all images deposited with this observation. Assistants imported information only from observations that showed at least one *Atta* reproductive, and recorded for each such observation the (a) date of observation; (b) date of submission to iNaturalist; (c) species of *Atta*, if known, for Research Grade records (otherwise record "unknown" if a species-ID was not known); (d) GPS coordinates; (e) locality information as displayed at iNaturalist under "Details" (town/city, state/province/district, country); (f) sex and wing-status of reproductives (alate female, dealate female, alate male; males are never dealate because they do not shed their wings, females shed their wings shortly after their mating flight); and (g) the URL of the iNaturalist record. Our primary dataset of observations of *Atta* reproductives deposited at iNaturalist covered a ten-year period from 2012 until the end of 2021, and including 20 countries across the entire *Atta* range from northern Argentina/Uruguay to the southern USA (Table S1, Data S1).

Because we were interested in characterizing the phenology of *Atta* mating flights on a fine geographic scale (i.e., by exact GPS location) and by ordinal (Julian) day in a year, we did not include records that had been submitted as "Private" at iNaturalist or did not provide the exact date of observation (some records deposited at iNaturalist obscure the exact observation date and list only the month of observation). We also excluded any images of pinned *Atta* reproductives in insect collections, because we were unsure whether the "observation" date submitted with such a specimen represented the collection date in the field, or a later date when a pinned specimen was photographed in an insect collection. Third, we excluded a few observations of dead specimens that were clearly dry, partly decayed, and the comment by the iNaturalist observer suggested that the specimen could have had its mating flight weeks or months before the observation date. Decisions regarding which observations to exclude or include in the final dataset, using the above criteria, were made blind (Kardish et al. 2015) with respect to any biogeographic patterns to be evaluated, and prior to any analyses of the final dataset (Data S1).

#### Data curation and verification

To minimize mistakes during data entry into our spreadsheet, we instructed assistants to double-check all information entered. One strategy to minimize mistakes was to enter, interpret, and curate only about 10 new records in a session, then take a break, and continue adding additional records to our spreadsheet only if still feeling concentrated enough to avoid mistakes. To eliminate any accidental mistakes and misidentifications, one of the coauthors (UGM) used the URLs entered in our Excel spreadsheet for each observation to compare the information recorded at iNaturalist with the information entered for each record in our Excel spreadsheet, verify all information, and rectify any mistakes. For example, determining the sex of an *Atta* alate was difficult in about 1% of the observations if the specimen was photographed from an angle that did not clearly show the head (the head of an *Atta* females is much larger than the head of an *Atta* male), or show the tip of the gaster with the partially everted genitalia characteristic of *Atta* males (females do not evert genitalia), or show the legs (male legs are thinner compared to the somewhat sturdier female legs). For observations where it was difficult to determine the sex, two coauthors (T.R.K. and U.G.M.) evaluated independently each of the respective images, discussed morphological features of a specimen to reach a consensus regarding its sex, or decided that sex could not be determined from a given image (these are recorded as "?" in Data S1).

#### Species identification

In the absence of corresponding workers, identifying *Atta* reproductives to species from external morphology is very difficult for both females and males (Borgmeier 1950, 1959). We therefore did not try to identify to species most iNaturalist observations of *Atta* reproductives included in our dataset, and we list these unidentified observations as "unknown" species (≈66% of all observations in our dataset cannot be identified to species; Table S1, Data S1). We list species names only if (i) a known expert naturalist had identified a given observation, or (ii) a single *Atta* species is known to occur in a given location. For example, only *Atta cephalotes* occurs in Belize and in Trinidad & Tobago; only *Atta mexicana* occurs in the Pacific States of Mexico; only *Atta* *cephalotes* occurs in north-east Costa Rica; only *Atta texana* occurs in Texas & Louisiana in the USA, and only *Atta mexicana* occurs in Arizona in the USA.

Because even expert taxonomists have great difficulty of identifying *Atta* reproductives to species (Borgmeier 1950, 1959), iNaturalist is actually the most comprehensive repository of observations of *Atta* reproductives, exceeding the number of *Atta* reproductives recorded at GBIF (Global Biodiversity Information Facility; https://www.gbif.org/). Specifically, the aforementioned ≈1500 observations in our dataset where an *Atta* reproductive could be identified unambiguously to genus at iNaturalist (but not to species level) are missing at GBIF, because GBIF uploads from iNaturalist only research-grade observations that have been identified to species by at least two naturalists. Consequently, far fewer records of *Atta* reproductives are deposited annually at GBIF than at iNaturalist. Although we do not know species identities for the majority of observations included in our dataset, it is likely that most or perhaps all of the currently 15 recognized *Atta* species (Bacci et al. 2009; Barrera et al. 2022) are represented in our dataset of 2335 observations, with the geographically more widespread species (*cephalotes*, *laevigata*, *mexicana*, *sexdens*) likely representing the majority in our dataset.

**Structure of the iNaturalist dataset**

In our dataset of 2335 observations, fewer observations were deposited at iNaturalist during the early years (Table S2) because of few users, but the number of records of *Atta* reproductives submitted per year increased gradually to 430 new records in 2019, then 743 and 812 new records for the Covid pandemic years 2020 and 2021, respectively, when more observers became active at iNaturalist (Table S2). About 67% of our total dataset were observations from only two years (2020 & 2021) and 85% of the total dataset were observations from the three years 2019-2021 (Table S2). Our aim was not to analyze differences in mating-flight patterns between years, and we therefore pooled all observations from 2012-2021 to estimate the phenology of mating flights in an average year during that time window.

Not surprising, countries with more iNaturalist observers (i.e., countries with more human inhabitants) contributed more observations to our total dataset (Table S1), a known pattern for observations uploaded to iNaturalist for any kind of organism (Di Cecco et al. 2021). Most observations were from locations in or near larger cities (Data S1), that is, areas with greater density of human populations, more users of iNaturalist, and consequently a greater likelihood of human-*Atta* encounters. Moreover, our impression was that the rate of observations tended to increase during weekends and on holidays. Unequal geographic and temporal coverage is typical for online community resources like iNaturalist (Di Cecco et al. 2021; Johnston et al. 2023). Two countries contributed disproportionally more observations, Colombia (n = 260 observations) and Mexico (n = 850 observations). These larger datasets from Colombia and Mexico allowed us to analyze the mating-flight patterns of *Atta* within each of these two countries on finer biogeographic scales (Figures 3-6).

**Observer biases**

The process of data accumulation at iNaturalist is not unbiased (Di Cecco et al. 2021). For example, we noticed that most *Atta* observations deposited at iNaturalist are from well-populated metropolitan areas or from countries with large human population sizes such as Mexico and Colombia, while observations from some countries were surprisingly underrepresented (e.g., Venezuela; Table S1). Frequencies of *Atta* observations at iNaturalist are actually combined measures of (i) the likelihood of human-*Atta* interaction (e.g., *Atta* reproductives can be attracted to lights in homes or to streetlights), and (ii) observer motivation to photograph the large, charismatic *Atta* reproductives and then upload images to iNaturalist. Geographic coverage of *Atta* at iNaturalist is therefore not uniform; some locations (e.g., cities, biological research stations) contribute disproportionately more observations, whereas rural locations are underrepresented. Such spatial heterogeneity in sampling effort is true also for many other biogeographic surveys (Johnston et al. 2022). In addition, observations at iNaturalist are biased across time, with disproportionately more observations reported on weekends, during holidays when more observers make time for outdoor activities, or during daytime rather than nighttime. It is therefore likely that observations of the *Atta* species flying at dawn or during the day are overrepresented in our dataset, whereas the *Atta* species flying late afternoon or at dusk (e.g., *vollenweideri*, *sexdens*) may be underrepresented because fewer observers collect observations after sunset. Because we did not analyze mating-flight patterns by individual *Atta* species (except for those locations in Mexico and Texas where only a single *Atta* species is known to occur, see above and Footnote 1 below), and because we pooled observations by month within a year (we pooled weekday and weekend observations within a month), we believe that any spatial and temporal observation biases do not distort our analyses of biogeographic patterns that we aim to elucidate.

Different observations at iNaturalist can be statistically dependent on each other (i.e., they are not independent). For example, observations of *Atta* reproductives are nested within mating-flight events triggered by local rainfall, and it is difficult to devise nested analyses that control for the effects of locality-specific rainfall patterns.

Second, different observers sometimes photographed and reported to iNaturalist the same individual *Atta* reproductive as different observation records, and we found even one case where an entire class of students reported to iNaturalist the same individual alate female, but this individual was photographed by different observers from different angles and deposited at iNaturalist as independent observations. Because these duplicate observations were submitted to iNaturalist at around the same time, we were able to recognize some such cases, and we therefore include for each such case only a single representative observation in our dataset (i.e., we excluded duplicates that we could recognize because of image similarities). It is possible that we did not detect all such duplicate submissions. Third, some observers clearly understood the difference between female and male *Atta* reproductives, and they reported to iNaturalist paired observations (an observation of a female and a separate observation of a male), but no observations of additional reproductives encountered at the same time. For these cases, we included in our dataset the observations of both the female and the male alate (Data S1), because we were also interested in evaluating approximate frequencies of female and male observations reported to iNaturalist.

**Validation and Ground-Truthing: Survey of Mating-Flight Observations by *Atta* Researchers**

To assess the validity of the biogeographic patterns inferred from our iNaturalist dataset, we contacted experts currently researching leafcutter-ant biology in locations across the entire *Atta* range. We asked each expert to (i) complete a questionnaire summarizing their personal observations on *Atta* mating flights (Table S3) and (ii) compare their own observations of *Atta* mating-flight phenology with the biogeographic patterns in mating-flight phenology apparent in the iNaturalist dataset (Figure 1). We emailed the questionnaire between Nov/2024-Nov/2025 to a total of 70 experts actively researching *Atta* biology; 48 experts returned a completed questionnaire (some groups of collaborating experts returned a single questionnaire); 8 experts declined and replied that they were too busy; and 16 experts did not respond to our initial email or a follow-up email later. All returned questionnaires are compiled in Table S3 in the order that questionnaires were received. The *Atta* researchers reported a total of 838 observations of *Atta* mating flights from across the entire *Atta* range and from the years 1975-2025 (Data S2), with most records from 2010-2025, approximately the same time window of the observations in our iNaturalist dataset (Data S1).

**Validation and Ground-Truthing: Survey of Mating-Flight Observations Reported in Literature**

To validate the biogeographic patterns inferred from our iNaturalist dataset (Figure 1), we completed an exhaustive review of published research articles, PhD dissertations, Master's theses, and web publications reporting any information on the time of year when *Atta* mating flights occur in specific locations, regions, or countries (Table S4). To find publications, we used Google Scholar and the search terms "Atta mating flight", "ant mating flight", "Atta nuptial flight", "Atta reproduction", "Atta reproductive", "Atta alate", "Atta vuelo nupcial", "Atta voo nupcial", "Atta vôo nupcial", as well as regionally-used names for *Atta* ants, including "saúva", "hormiga arriera", "hormiga forrajera", "zompopo", "zompopo de mayo", "zángano", "wiwi", "bibijagua", "bachac", "cuqui", "coqui", "siquizapa", "cushi", "acoushi", "coushi", "cutter ant", and "parasol ant"; then used the Google-Scholar functions "Cited by" and "Related articles" associated with each *Atta* publication to find additional publications. We obtained PDFs for all publications listed in Table S4 directly from Google Scholar or through the University of Texas Library (https://lib.utexas.edu/). We ended our literature search in December 2025, so publications post-dating 2025 are not included in our survey. Our literature survey found 806 records of *Atta* mating flights from across the entire *Atta* range and from the years 1894-2025 (Data S3; Table S4), with most records from the years 1940-2025.

**Expanded dataset for *Atta* in north-east Mexico and Texas**

While analyzing latitudinal patterns of mating-flight phenology of *A. texana* in Texas and Louisiana (see preceding paragraph), we noticed a marked latitudinal switch from mating flights predominantly in August-October in southmost Texas to March-May in all other latitudes across Texas. To understand this shift within a greater latitudinal context, we extended our latitudinal transect across Texas southward to include observations from *Atta* populations in the lowlands along the Gulf of Mexico as far south as northern Veracruz State. This transect spans an area of sympatry between *A. texana* and *A. mexicana* in the States of Tamaulipas and Nuevo Leon in the north-east border region of Mexico (only *A. texana* exists north of that border region in Mexico, only *A. mexicana* exits south of that border region in Mexico). We restricted this transect to latitudes north of latitude N20° near Xalapa (i.e., we excluded the southern half of the State of Veracruz) because somewhat south of latitude N20°, *A. mexicana* is sympatric with *A. cephalotes*. The entire 1500-km transect ranges from latitude N20° in Veracruz State across Tamaulipas and Texas to N33° in northern Texas. For the region in Mexico covered by this transect, we included only records from the Gulf Coast lowlands (i.e., we excluded records from the mountainous areas to the west of the Gulf Coast lowlands in Mexico), because *Atta* mating flights occur slightly later in the year in the more western mountainous areas than in the Gulf Coast lowlands.

**Rainfall As a Trigger of Mating Flights in Different Ecoregions of Colombia**

In addition to GPS information, iNaturalist observations are structured for each country into administrative units (e.g., City/Municipio, State/Estado/Provincia/Departamento), and we transcribed this information into our primary dataset (Data S1) to group observations for some of our biogeographic analyses (e.g., Figure S2). Because a Departamento (Department) in Colombia can span different ecoregions, we characterized annual mating-flight phenology also by ecoregions as defined by Dinerstein et al. (2017). To analyze relationships between mating-flight and rainfall patterns in different ecoregions in Colombia, we used the rainfall dataset from Urrea et al. (2019) (replicated in Data S5), in which rainfall patterns are defined by up to four components that describe rainfall periods of increased precipitation (referred to as "seasons"). These components capture bimodal or multimodal rain patterns typical of tropical Andean climates. Each "season" is defined by a start day of the year (SOS), duration in days (DOS), and total rainfall (mm) in that season. To estimate daily rainfall, we distributed the total rainfall of each season uniformly across its days of duration (total rainfall/DOS), and we summed the contribution for all four seasons for each day of the year. Mating flights and rainfall datasets were processed, standardized, and paired by day of the year (DOY) (Data S5). We grouped both rainfall and mating flight observations by ecoregions with similar climatic conditions in Colombia. To enable visualization in the same graph, we normalized (z-scored) both mating-flight observations and daily rainfall values, so both normalized mating-flight and rainfall values can be projected onto the same y-axis of a single graph. We quantified the temporal relationship between rainfall and mating flights by computing lag-0 correlations, followed by cross-correlation analyses over a ±30-day window to identify the lag with the highest correlation, using the methods of Curriero et al. (2005). We evaluated the correlation between mating flight patterns and rainfall using RStudio version 4.6.0 (2026) (R-Scripts S1 and S2).

**Rainfall Patterns in 2012-2021 Corresponding to iNaturalist Records**

To understand how regional rainfall patterns drive *Atta* mating-flight patterns, we obtained monthly total surface precipitation data from the *Precipitation 1.0 degree Data Full Reanalysis* dataset available from the Global Precipitation Climatology Centre (GPCC; https://psl.noaa.gov/data/gridded/data.gpcc.html; Schneider et al. 2022) for the ten years 2012-2021 covered also by our iNaturalist dataset. We imported mean monthly precipitation data available at GPCC into ESRI ArcMaps10.8, projected data in WGS1984, and smoothed data using bilinear interpolation. To draw rainfall maps, we acquired country outline and lake outline shapefiles from EfrainMaps (http://www.efrainmaps.es; Tapiquén 2020), and ocean shapefiles from NaturalEarth (https://www.naturalearthdata.com). Figure S1 shows the 120 maps (12 months X 10 years) of these monthly rainfall patterns across the *Atta* range from South to North America.

From these monthly rainfall patterns, we inferred maps summarizing local rainfall averages for each month across the ten years 2012-2021 (Figure 2). We calculated monthly means across the 10-year window using cell statistics in the spatial analyst toolbox and we smoothed the resultant rasters using bilinear interpolation. The average rainfall visualized in Figure 2 allowed us to derive expectations regarding corresponding changes in *Atta* mating-flight patterns. For example, in the month-by-month series of annual rainfall patterns across a transect from Central America to North America (Figure 2), we noticed that, following a drier period in January-March during an average year, the wet season start earliest in eastern Panamá in March/April, then the onset of the wet season spread north-westward across Central America in May, and then north-westward across Mexico in June/July (Figure 2). These annual shifts in local onset of the wet season predict corresponding shifts in the annual local onset of *Atta* mating flights ranging from April in Panamá to July in north-west Mexico.

**FOOTNOTES**

**Footnote 1:**
Most observations of *Atta* reproductives at iNaturalist are identified only to genus level (Data S1); for potential between-species comparisons, sample sizes of *Atta* reproductives identified to *Atta* species at iNaturalist are therefore very small (Table S1). For example, in the iNaturalist dataset from north-west South America, only 4.9% of the observations of *Atta* reproductives are identified to species (14 *Atta cephalotes* and 3 *Atta laevigata* observations; the remaining 342 observations are unidentified *Atta* species; Table S1). Such sample sizes are insufficient to test whether possible species-specific mating-flight specializations on different seasons contribute to an overall annual mating-flight pattern in a region, like the annual bimodal mating-flight phenology in Figure 1 D for north-west South America. The lack of sufficient sample sizes of identified *Atta* species therefore precludes robust species comparisons using iNaturalist data, except for *A. mexicana* and *A. texana* in north-east Mexico and south-central USA (Section 3.6 and Figure 8 in main article).

Whereas the great majority of observations in our iNaturalist dataset are identified only to genus *Atta* (few observations are identified to species), many observations of *Atta* reproductives reported in the questionnaire survey (Table S3, Data S2) and in the literature survey (Table S4, Data S3) are identified to *Atta* species. We combined these data from the questionnaire and literature surveys into a single dataset (Table S5), to explore whether sympatric *Atta* species may differ in mating-flight phenology in a specific region or in a specific country (Table S5). Preliminary analyses (Table S5) did not find clear differences in mating-flight phenology between sympatric *Atta* species, except for the aforementioned difference between *A. mexicana* and *A. texana* across the contact zone of these two species in north-east Mexico (Figure 8 in main article). Within all other regions or countries, either sample sizes of individual *Atta* species were too small to permit robust comparisons between sympatric *Atta* species (Table S5); or, for those *Atta* species with sufficient sample sizes, mating-flight phenologies do not differ between sympatric *Atta* species (Table S5). That is, the available data (Table S5) suggest that sympatric *Atta* species have similar or identical mating-flight phenologies.

Elucidating differences in reproductive life-history between different *Atta* species was not an aim of the present study, and we therefore did not explore species differences beyond the preliminary analyses in Table S5. We will elucidate between-species differences in greater detail in a separate study focused on differences in time-of-day of mating flights (dawn, morning, afternoon, dusk) between the 15 recognized *Atta* species (Mueller et al. in preparation).

**Footnote 2:**
The blue dashed line in Figure 1 in the main article was inferred entirely from our iNaturalist records and was drawn blindly without knowledge of prior publications. The blue dashed line in our Figure 1 approximates a demarcation in *Atta* mating-flight phenology between southern and northern Brazil highlighted in Forti and Boaretto (1997). [In contrast, our inferred mating-flight phenology in southern Brazil (Figure 1) differs from the one shown by Forti and Boaretto for southern Brazil (FIGURA 7 shown at right)]. We have added to FIGURA 7 of Forti and Boaretto (1997) a dashed red line approximating the dashed blue line shown in our Figure 1. Forti and Boaretto (1997) and our analyses (Figure 1) highlight the distinct mating-flight phenologies between northern Brazil and the rest of Brazil.

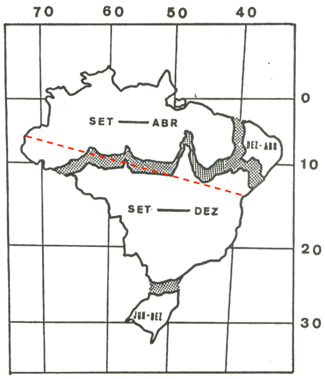

FIGURA 7 - Épocas de ocorrência de vôo nupcial para as saúvas do Brasil, em diferentes regiões.

The dashed blue line in our Figure 1 is drawn across South America from southernmost Ecuador at the Pacific coast to the northern border of the State of Minas Gerais in Brazil near the Atlantic coast (for the Brazil portion of this line, the line is approximated by the dashed red line added here to FIGURA 7 of Forti and Boaretto 1997). We regard the dashed blue line in Figure 1 as a preliminary demarcation, which we drew after inspecting our iNaturalist dataset, then adjusting the line such that, south of that line, iNaturalist observations of *Atta* reproductives occurred almost exclusively in the austral spring (September to December; Figure 1H). That is, we did not conduct a rigorous discriminatory analysis to determine transition zones in mating-flight phenology across South America. We see the blue line in Figure 1 as a starting point to explore biogeographic patterns more rigorously in future analyses, using ideally an expanded dataset that also incorporates iNaturalist observations collected after 2021 (our dataset covers 2012-2021). It is possible that *Atta* mating flight patterns shift in northern and central South America depending on cyclical changes in rainfall, like those driven by temperature changes in the Pacific Ocean and south Atlantic Ocean (Yoon and Zeng 2010; Marengo and Espinoza 2016).

**Footnote 3:**
Previous reports noted the difference in *Atta* flight season between north-east and southern Brazil (Rêgo and Brandão Filho 1941; Forti and Boaretto 1997; Autuori 2010; Santos-Maríno et al. 2011; Jaccoud 2017), and some reports refer to an extended *Atta* flight season of half a year from November to April in north-east Brazil (Rêgo and Brandão Filho 1941; Delabie 2002, Santos-Maríno et al. 2011; Jaccoud 2017) or mention the possibility of *Atta* flights departing from single nests at two distinct times per year in Bahia ("Dêstes formigueiros … duas vêzes por ano saem enxames de içás ou tanajuras"; page 7, Gonçalves 1951). Our sparse records from iNaturalist suggest possibly shorter flight seasons in the Brazilian States of Ceará (December and January, n=9 records) and Pernambuco (February to April, n=5 records), and possibly an extended 6-month flight season from November to April in the State of Bahia (n=13 records), but it is premature to draw conclusions from our limited dataset of only 50 observations for all of north-east South America (i.e., north-east Brazil).

**Footnote 4:**
In a study by Gejskes (1953) in Suriname in north-central South America, Gejskes excavated *Atta* nests monthly throughout the year, and he noted the limited time of the year when he found reproductive brood in *Atta* nests. Reproductive brood develops in nests for several months preceding the actual mating-flight season, and Gejskes found this period of reproductive brood production to be November-June for *A. cephalotes*, and October-April for *A. sexdens*. This suggested to Gejskes "a slight difference between the two [*Atta* species] concerning their breeding time" (page 182). Further, Geijskes illustrated graphically (page 183) a "small rainy season" in December and January when he believed *A. sexdens* has mating flights, and a "main rainy season" in April to June when he believed *A. cephalotes* has mating flights. While this hypothesized separation by four months of mating flights between the two *Atta* species is intriguing, we have found in our field work that, in any given locality, all sympatric *Atta* species tend to swarm in the same months and in at least some locations on the same day triggered locally by the same preceding rain, but different *Atta* species are reproductively isolated because they swarm at different times of the day (e.g., Questionnaires #01, #05, #16, #17, #18, #23, #30). See also the preliminary comparisons in Table S5 showing no indication that different sympatric *Atta* species have mating flights in different seasons of the year.

**Footnote 5:**
The predominance of *Atta* mating flights in May throughout Central America (Figures 5B-D, main article) explains why *Atta* leafcutter ants are called "hormigas de Mayo" (May ants) in Costa Rica and Panamá, or "zompopos de Mayo" in north-west Central America (in one of the Mayan languages, "zonm" apparently means ant, and "popo" means big; Luna 2021). In contrast, the predominance of *Atta* mating flights in April in north-west Colombia (Figure 5A) and the predominance of *Atta* mating flights in June and July in Mexico (Figures 5E-O) explains why *Atta* are *not* called "hormigas de Mayo" in Colombia and Mexico. That is, *Atta* are called "hormigas de Mayo" only in Central America, but not outside of Central America, matching the geographically-varying mating-flight phenologies shown in Figures 5A-O.

**Footnote 6:**
An interesting feature in Figures 6A&B is that, while most of the observations are concentrated for each latitudinal or longitudinal bracket within a narrow time window corresponding to the onset of spring rains at a given latitude or longitude, there exist also a few very early observations in winter well before the onset of spring rains (see outlier black dots towards bottom of Figures 6A&B). The reasons for these outlier observations could be (a) observers submitted to iNaturalist records with incorrect dates; or (b) chance departures of a few reproductives from *Atta* nests in winter, perhaps stimulated by minor rains or other factors. The absence of such early records at higher latitudes where it is too cold in winter for reproductives to fly (rightmost latitudinal brackets in Figure 6A), suggests that these unusually early flight records may be real, and are not entirely due to errors (if the outliers were all submission errors, such errors would presumably also affect some observations at higher latitudes).

**Footnote 7:**
The late-year mating-flight phenology of *A. texana* reproductives from latitudes N26-27° in Texas is unique within the range of *A. texana* and different from the spring mating-flight phenology of *A. texana* reproductives from higher latitudes N27-33° in Texas (Figure 8 in main article). This season switch at around N27° latitude from early-year to late-year mating flights appears to be real and cannot be attributed to a sample-size artifact, because (a) the number of observations for latitudes N26-27° seem sufficient (n=27) to characterize of mating-flight phenology, and (b) specialization on late mating flights at latitude N26-27° is consistent across each of the seven years 2018-2024 for which observations are available at iNaturalist (only late-flying *Atta* and no early-

flying *Atta* were observed in each of these seven years for the latitudinal bracket N26-27°; no records are available at iNaturalist from before 2018 for this latitudinal bracket). Observer bias (e.g., absence of observers in spring) is not an explanation for this late-flying pattern, because iNaturalist observations of other ant species and other organisms from southmost Texas are mostly from spring, when many naturalists visit this area because of mild spring temperatures. Only *A. texana* is known from southmost Texas (e.g., UGM examined over 100 *Atta* nests in the latitudinal bracket N26-27° in Texas, and found only *A. texana* there), making it unlikely that all the observed late-flying *Atta* (n=27) there are *A. mexicana* dispersing into Texas from populations in Mexico. Lastly, rainfall or temperature patterns between latitudinal brackets N26-27° and N27-28° are not radically different, suggesting that other factors drive a switch from early-flying to late-flying in southmost Texas. These factors are further discussed in Section 3.6 and Figure 8 in the main article, placing the mating-flight phenologies of *A. texana* in the context of a latitudinal transect covering also *A. mexicana* and ranging from N20° latitude in Mexico to the northernmost *Atta* populations at latitude N33°.

**Footnote 8:**
The 1500-kilometer transect in Figure 8 starts at 20° latitude in the northern half of the State of Veracruz in Mexico and continues northward along the Gulf Coast, to include across this range only records from *A. mexicana* from the Gulf Coast lowlands of Mexico. Somewhere south of latitude 20° in Mexico somewhat south of Xalapa (latitude 19.5), the range of *A. mexicana* starts to overlap with the range of *A. cephalotes* (García-Martínez et al 2015; Baena et al 2020; Gómez-Díaz et al 2023), and we therefore excluded from the 1500-kilometer transect *Atta* records south of 20° latitude, because our aim was to analyze a transect including only the two species *A. mexicana* and *A. texana*. *A. mexicana* has so far not been found in Texas in ant surveys (Wheeler and Wheeler 1985; O'Keefe et al. 2000; UGM unpublished observation). One of us (UGM) has searched extensively for *A. mexicana* in southmost Texas and in the entire border region in the USA along the Rio Grande River from the Gulf of Mexico to Del Rio (westernmost *Atta* populations in Texas) (Figure 8), and all the 250+ nests from that region for which soldiers could be examined were identified as *A. texana* (soldiers of *A. mexicana* have a distinctive shiny head and gaster, whereas soldiers of *A. texana* have a non-shiny matte head and gaster). *A. mexicana* therefore does not occur in the Texas, or perhaps at such low frequency that the presence of *A. mexicana* in Texas has so far not been recorded.

*A. mexicana* and *A. texana* are sympatric in north-east Mexico in the States of eastern Coahuila, northern Nuevo León (Figure 8H), and northern Tamaulipas. Unfortunately, no records of *Atta* reproductives are available at iNaturalist for that biogeographically interesting region in northern Tamaulipas, and other surveys (e.g., Alatorre-Bracamontes et al. 2010; Flores-Maldonado et al. 2021) documented a paucity of *Atta* observations for this region (see Figure 1 on page 83 in Flores-Maldonado et al. 2021), most likely because field biologists avoid the unsafe region just south of the Mexico-US border. Only Sánchez-Peña (2005) conducted a survey specifically aimed at *Atta* in this border region of Mexico. Both *A. mexicana* and *A. texana* appear very rare in northern Tamaulipas (Sánchez-Peña 2005) and populations seem to be fragmented in north-eastern Mexico (Sánchez-Peña 2010), most likely because of local lack of moisture, intense agriculture, agricultural irrigation causing *Atta* nests to drown, and extensive pesticide use. Sánchez-Peña surveyed the entire area in July and August 2000 and found the northernmost populations of *A. mexicana* near San Fernando in Tamaulipas, in the Monterrey area in Nuevo León, in Sabinas Hidalgo in Nuevo León, and in Sabinas Coahuila in Coahuila. Sánchez-Peña found only *A. mexicana*, but not *A. texana* at each of these locations, and he did not find any *Atta* nests between these locations and the Rio Grande River (Figure 2.1 in Sánchez-Peña 2005), confirming for Nuevo León essentially the same northern range limit of *A. mexicana* found in an earlier ant survey (Rodríguez Garza 1986). Somewhat further south from this currently predicted northern range-limit of *A. mexicana*, Sánchez-Peña found that "*A. mexicana* was widespread between Pesquería and Monterrey, in Nuevo León; between Monterrey and Ciudad Victoria, Tamaulipas; in Tamaulipas, between Ciudad Victoria and Soto La Marina, and between Soto La Marina and San Fernando. Along the Gulf of Mexico coast of Tamaulipas, and traveling on a south-north direction, the distribution of *A. mexicana* appeared to be continuous all the way to the city of San Fernando, where it was observed about 100 km south of the Texas border" (pages 7&8, Sánchez-Peña 2005). This suggested to Sánchez-Peña (2005) that any sympatry between *A. mexicana* and *A. texana* is limited to a zone ranging from eastern Coahuila (Parque Rio Sabinas, 13 km north-west of Melchor Múzquiz; Sánchez-Peña 2010) through northern Nuevo Leon and northern Tamaulipas to the Gulf of Mexico. We highlight this band of most likely sympatry between *A. mexicana* and *A. texana* in Figure 8.
